# Asymmetric learning and adaptability to changes in relational structure during transitive inference

**DOI:** 10.1101/2024.07.03.601844

**Authors:** Thomas A. Graham, Bernhard Spitzer

**Affiliations:** Research Group Adaptive Memory and Decision Making, Max Planck Institute for Human Development, Lentzeallee 94, 14195 Berlin, Germany; Max Planck Institute for Human Cognitive and Brain Sciences, Stephanstraße 1A, 04103 Leipzig, Germany; Max Planck School of Cognition, Stephanstraße 1A, 04103 Leipzig, Germany

**Author notes:** Corresponding author: Thomas A. Graham.

## Abstract

Humans and other animals can generalise from local to global relationships in a transitive manner. Recent research has shown that asymmetrically biased learning, where beliefs about only the winners (or losers) of local comparisons are updated, is well-suited for inferring relational structures from sparse feedback. However, less is known about how belief-updating biases intersect with humans’ capacity to adapt to changes in relational structure, where re-valuing an item may have downstream implications for inferential knowledge pertaining to unchanged items. We designed a transitive inference paradigm involving one of two possible changepoints for which an asymmetric (winner-or loser-biased) learning policy was more or less optimal. Participants (N=83) exhibited differential sensitivity to changes in relational structure: whereas participants readily learned that a hitherto low-ranking item increased its rank, moving a high-ranking item down the hierarchy impaired downstream inferential knowledge. Behaviour best captured by an adaptive reinforcement learning model which exhibited a predominantly winner-biased learning policy but also modulated its degree of asymmetry as a function of its choice preference strength. Our results indicate that asymmetric learning not only accounts for efficient inference of latent relational structures, but also for differences in the ease with which learners accommodate structural changes.

**Author Summary:** When reasoning about relationships between objects, events, or people, humans can readily use previous experiences to infer relations that they have never encountered before. For example, if Anna beats Bruce at tennis, and Bruce beats Clara, then one can predict that Anna will likely also beat Clara. Human learning in such ‘transitive inference’ problems tends to be winner-biased – that is, upon observing Anna’s victory over Bruce, a spectator would be more likely to attribute this outcome to Anna’s skill than to Bruce’s lack thereof. However, in a constantly changing world whose comparative relations are rarely static, humans must also be able to infer how changes in the outcomes of certain comparisons bear on other relationships within a transitive hierarchy. Combining behavioural testing and computational modelling, we show that a learning strategy that preferentially focuses on the winners of comparisons induces greater flexibility for certain types of hierarchy changes than for others. In addition, we provide evidence that humans may dynamically adjust their degree of learning asymmetry according to the current strength of their beliefs about the relations under comparison.

## Introduction

Humans readily learn how items rank on a variety of latent scales, such as those pertaining to hedonic or economic value, or social influence. Such representations of rank permit novel inferences of indirectly related states or entities. For instance, knowing that A<B and B<C enables one to infer, through transitive inference (TI), that A<C. TI has been widely studied in humans, non-human primates, rats and birds alike [1–4]. Under TI learning regimes, training trials offer participants trial-and-error feedback about pairwise comparisons between items of neighbouring rank, which must then be used to infer unseen test relations between non-neighbouring items. In requiring agents to use the outcomes of pairwise comparisons to update their estimates of the rankings within a linear set, TI paradigms lend themselves to the application of simple reinforcement learning (RL) frameworks that model the influence of choice feedback on the subjective value of the compared items. Recent work adopting this approach demonstrated that TI learning is characterised by, and indeed benefits from, an asymmetric policy under which either the winner (or the loser) of a pair is preferentially updated [5]. Specifically, this benefit emerged in simple RL models furnished with separable, or ‘asymmetric’ learning rates for updating winners and losers, with most participants displaying a bias towards updating winners. This cognitive distortion during inferential learning fits into a wider body of literature on human biases towards positive [6,7] or confirmatory feedback signals [8– 11].

The constantly changing nature of an agent’s environment necessitates that any capacity for relational learning must exhibit adaptability, while also ensuring robustness [12]. The learning dynamics underlying humans’ ability to adapt to volatile reward environments have been studied in tasks involving changepoints or reversals [13–15]. Likewise, sensory preconditioning paradigms have been used to investigate the conditions under which relational representations are retrospectively re-evaluated via relearning associations between rewarded and indirectly related stimuli, or through inference at the time of choice [16,17]. These studies have demonstrated humans’ ability to infer how changes in local reward feedback pertain to indirectly related stimuli, underscoring the utility of changepoint manipulations in probing inferential learning capabilities.

Studying changepoints in larger relational structures allows one to investigate how agents rapidly modify existing knowledge in response to minimal new information [2,18]. Less is known, however, about how such ‘few-shot’ local relational changes impact downstream inferential knowledge, nor how this capacity to adapt to changes in relational structure intersects with well-documented belief-updating biases in humans. Consider a sports league where a spectator learns how the teams rank with respect to one another based on the outcomes of head-to-head matches between them. Halfway through the season, the unexpected loss of the reigning champions against a team sitting at the bottom of the hierarchy may be indicative of the former’s fall from grace, and/or the latter’s resurgence. Ascertaining which team’s ranking has changed will thus determine how much one needs to update one’s predictions about how this team will fare against others in the league, while ensuring minimal disruption to knowledge pertaining to the relations between teams whose performance remains unchanged (Fig 1A). Interestingly, a corollary of the asymmetric RL framework is that the ease with which such changes in relational structure are accommodated, and thus any resultant impact on downstream inferential knowledge, should vary as a function of the asymmetry in an agent’s learning policy (see Fig 1C and S1 for simulations). If humans are biased towards preferentially increasing their estimates of winners, then the sudden decline of the hitherto best team to the bottom of the leaderboard should be less readily accommodated than the rapid ascendency of the worst team to the top of the table. The relative difficulty with which this former change in ground truth structure is learned would also, in turn, reduce the discriminability of mid-table teams whose rankings remain unchanged, and thus disrupt the agent’s inferential knowledge with respect to the middle of the transitive hierarchy.

**Fig 1.**
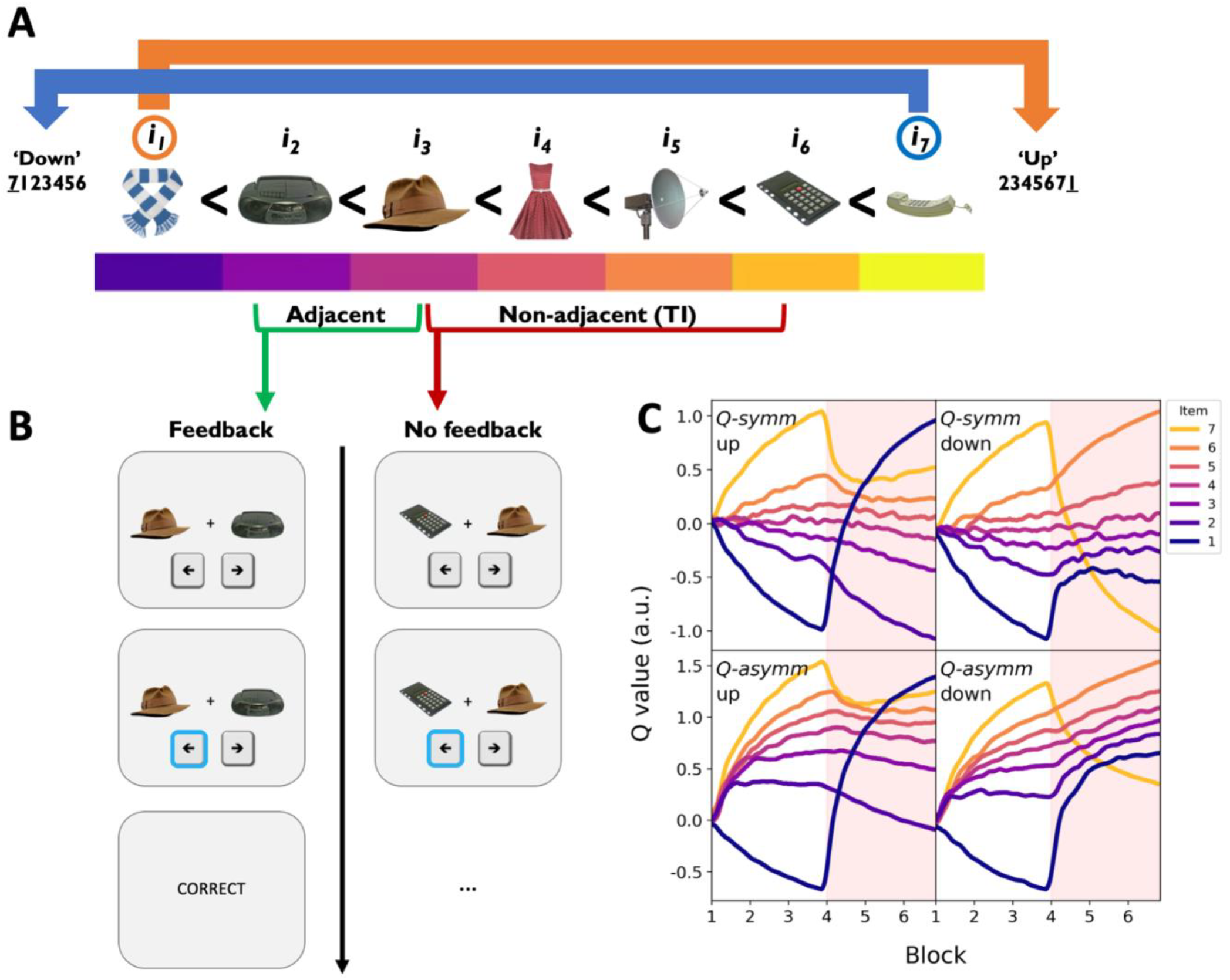
Experimental paradigm and model simulations. **A)** Example ‘cnarciness’ rankings of a set of seven items in an ordinal hierarchy. After three blocks, the ground truth structure changed in one of two possible ways: in the ‘down’ group (blue), the most cnarcy item *i7* (here, the telephone) moved to the bottom of the hierarchy, whereas in the ‘up’ group (orange), the least cnarcy item *i1* (here, the scarf) moved to the top of the hierarchy. **B)** On each trial, participants were asked to choose which of two items they believed to be the most cnarcy. Binary feedback was delivered on adjacent trials containing items neighbouring in rank (green), while TI comparisons between non-adjacent items offered no feedback (red). **C)** Simulated item value estimates (‘Q values’) under the symmetric agent *Q-symm* (top row) and the asymmetric agent *Q-asymm* (bottom row) for the ‘up’ and ‘down’ experimental conditions (left and right columns, respectively). Red shaded half of each panel represents the post-changepoint phase of the experiment. Whereas non-anchor item value estimates are equally discriminable following both changepoints under *Q-symm’s* symmetric learning policy, *Q-asymm* predicts impaired discriminability of item values in the ‘down’ condition relative to the ‘up’ condition (cf. Fig S1). Models were simulated using parameter ranges consistent with participant learning asymmetries reported by Ciranka et al. [5].

Accordingly, there is evidence to suggest that the preferential integration of positive reward prediction errors can lead to choice inertia when the best and worst options in a two-armed bandit are flipped [19–21]. Likewise, humans are more reluctant to revise their subjective beliefs about the quality of a deteriorating foraging environment, relative to an environment whose reward rate improves [22]. While these studies support the idea that positively biased agents are more sensitive to positive changes in the value of options and reward environments, the prediction that the biased reorganisation of relational knowledge should have a downstream impact on unchanged elements of a transitive hierarchy remains untested. Moreover, while these predictions are made under the assumption of a static degree of learning asymmetry, introducing a changepoint in a TI learning paradigm also allows one to explore whether learning asymmetries may dynamically adjust or even reverse in a task-dependent manner, a possibility for which empirical evidence in other learning regimes is mixed [23,24; but see 25].

Here, we therefore sought to investigate whether biased learning policies confer different levels of (in-)flexibility to changes in an environment’s relational structure. Participants (N=83) performed a TI paradigm involving one of two possible changepoints for which a winner-biased learning policy was more or less optimal. In addition to replicating previously observed learning asymmetries in the pre-changepoint task phase, we found evidence supporting our model prediction that such biased learning strategies differentially advantage agents’ ability to accommodate directional shifts in the environment’s underlying relational structure. Computational modelling of behaviour further revealed that such differential sensitivity was best captured by an extension of our asymmetric RL model whose degree of learning rate asymmetry varied as a function of the strength of its choice preference. We thus provide a parsimonious account for how learning rate asymmetries may dynamically adapt to task conditions, unifying our present findings with previous research into belief-updating biases.

## Results

### Changepoint TI Paradigm

Participants (N=83) performed a computerised task in which they were, on each trial, presented with two items drawn from a set of seven *i*_*1*_, *i*_*2*_,*… i*_*7*_, and instructed to choose the item that they thought was more ‘cnarcy’ than the other using a button press. The relative cnarciness of each item was established at the beginning of the experiment by randomly assigning a ground truth rank from 1-7 to each item, such that *i*_*1*_ and *i*_*7*_ represented the least and most cnarcy items respectively. On ‘adjacent’ trials comparing items with neighbouring ranks, participants received deterministic feedback about whether they had correctly/incorrectly chosen the more cnarcy item. In contrast, on ‘TI’ trials comparing non-neighbour items, participants did not receive any feedback. Thus, participants were required to use sparse feedback from pairwise comparisons between adjacently ranked items to infer the transitive hierarchy governing the item set (Fig 1A-B).

Adjacent and TI trials were randomly interleaved within each of six blocks, allowing us to examine the evolution of TI over time. Critically, after the third block, a minimal change in the items’ hierarchy was introduced: in the ‘up’ group of participants (N=39), the hitherto lowest-ranking item *i*_*1*_ moved up the hierarchy to become the highest-ranking item, while in the ‘down’ group (N=44), the highest-ranking item *i*_*7*_ moved down to become the lowest-ranking item. In both groups, the relations between all other items remained exactly as they were before, such that the new ranking of item-IDs from lowest to highest could be represented as *7123456* in the ‘down’ group, and *2345671* in the ‘up’ group. Since participants only received choice feedback for adjacently ranked items, this change in the underlying ground truth only resulted in minor changes in the feedback received by each group. Specifically, on trials comparing the newly adjacent items *i*_*1*_ and *i*_*7*_, participants in both groups received new feedback consistent with *i*_*7*_<*i*_*1*_. The only difference between the two groups was in the two comparisons for which feedback was *removed* as a result of the rank change: ‘down’ participants no longer received feedback on trials comparing *i*_*6*_ *vs. i*_*7*_, whereas ‘up’ participants no longer received feedback on trials comparing *i*_*1*_ vs. *i*_*2*_, since these pairs of items were no longer adjacently ranked in each case. Thus, the objective changes in the underlying hierarchy could only possibly be inferred on the basis of two pieces of information: 1) the newly introduced *i7*<*i*_*1*_ relation, 2) the persistence or omission of the *i*_*1*_<*i*_*2*_ or *i*_*6*_<*i*_*7*_ relation.

### Simulations

Following Ciranka et al. [5], we simulated relational learning in our TI paradigm using simple RL models that updated the value (i.e. ‘cnarciness’) estimates *Q* of winning and losing items *x* and *y*, respectively, following choice feedback under a modified Rescorla-Wagner updating rule [26]:

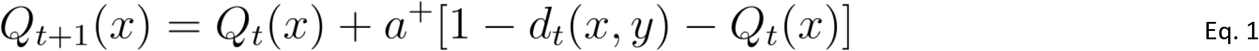

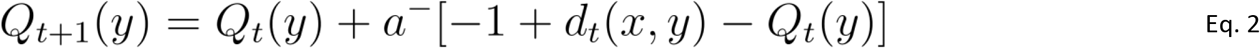

where *a*^*+*^ and *a*^*-*^ are the learning rates for winners and losers respectively. Separating these learning rates allowed the model to implement varying degrees of symmetry/asymmetry in its learning policy. We defined the symmetric model *Q-symm* as an agent for whom *a*^*+*^ = *a*^*-*^, meaning the agent increased and decreased its value estimates for winners and losers of each choice outcome respectively by equal amounts. In contrast, we defined the asymmetric model *Q-asymm* as an agent whose learning rates *a*^*+*^ and *a*^*-*^ could freely vary. In the case where *a*^*+*^ > *a*^*-*^, the agent was ‘winner-biased’, disproportionately increasing its value estimate for a comparison’s winner relative to its loser, whereas the agent was ‘loser-biased’ if *a*^*+*^ < *a*^*-*^.

Value updates were scaled by the relative difference between *Q*_*t*_*(x)* and *Q*_*t*_*(y)*, as represented by the *d*_*t*_*(x,y)* term in Eqs. 1-2 (see Eq. 4 in *Materials and Methods*, ‘Behavioural Models’). We modelled the probability of choosing *i*_*x*_>*i*_*y*_ as a sigmoid function of the difference between the estimated item values, scaled by a noise or ‘temperature’ parameter *τ* (see Eq. 5 in *Materials and Methods*, ‘Behavioural Models’).

We first present simulations of the symmetric and asymmetric RL agents *Q-symm* and *Q-asymm*, respectively, in order to derive model-based predictions for how humans should behave in our changepoint TI paradigm (Fig 1C and S1). We simulated model performance over a range of parameter values matching those previously estimated to fit human TI behaviour by Ciranka et al. [5], where participants tended to exhibit a winner-biased learning policy (i.e. *a*^*+*^ > *a*^*-*^) when fitted with *Q-asymm*. Preferentially updating winners in this way leads to compression of *Q-asymm’s* latent value structure before the changepoint, such that pairs of higher valued items are less discriminable than lower valued items. This reduced sensitivity towards larger values is a signature of asymmetry in relational learning. In contrast, the symmetric agent *Q-symm* exhibits no such compression (for details, see [5]).

Interestingly, *Q-asymm’s* asymmetric learning policy predicts a difference in how efficiently it should adapt to our changepoint manipulation in the ‘up’ condition relative to the ‘down’ condition (Fig 1C and S1). If learning is biased towards winners, the changepoint in the ‘up’ condition should be easily accommodated, since *Q-asymm* selectively and appropriately increases its value estimate for *i*_*1*_ without needing to update any other items. On the other hand, in the ‘down’ condition, *Q-asymm’s* initial tendency to increase its estimate for *i*_*1*_ over-inflates this item’s value, and underestimates *i*_*7*_’s decline in value. In contrast, *Q-symm’s* proportionate updating of winners and losers means that it will adapt to these two objective changes in the underlying ground truth with equal efficiency. Thus, if inferential learning is characterised by an asymmetric, winner-biased learning policy, then this yields the empirical prediction that humans should more efficiently adapt to the change in relational structure in the ‘up’ condition than in the ‘down’ condition.

### Value Compression

Focusing first on participants’ pre-changepoint behaviour (that is, all trials preceding the first *i*_*7*_*<i*_*1*_ trial in the fourth block), we confirmed that participants not only learned the cnarciness relations between items of neighbouring rank, but also used the feedback from these trials to accomplish TI (Fig 2, leftmost column). Participants in both groups exhibited above-chance accuracy both on pre-changepoint trials involving adjacent items (‘up’: mean accuracy = 0.67 ± 0.01 SE, *t*(38) = 12.94, *p* < .001; ‘down’: mean accuracy = 0.65 ± 0.01 SE, *t*(43) = 10.35, *p* < .001), and on pre-changepoint TI trials (‘up’: mean accuracy = 0.73 ± 0.02 SE, *t*(38) = 13.22, *p* < .001; ‘down’: mean accuracy = 0.75 ± 0.01 SE, *t*(43) = 16.67, *p* < .001). In both groups, we also found evidence for the widely observed ‘symbolic distance effect’ [27,28] in both pre-changepoint accuracy and reaction time (RT) data, such that greater ordinal distance between comparanda on TI trials was associated with higher accuracy (‘up’: *β* = 0.04, *t*(38) = 8.30, *p* < .001; ‘down’: *β* = 0.05, *t*(43) = 11.07, *p* < .001) and faster responses (‘up’: *β* = -0.03, *t*(38) = -4.11, *p* < .001; ‘down’: *β* = -0.03, *t*(43) = -5.24, *p* < .001).

**Fig 2.**
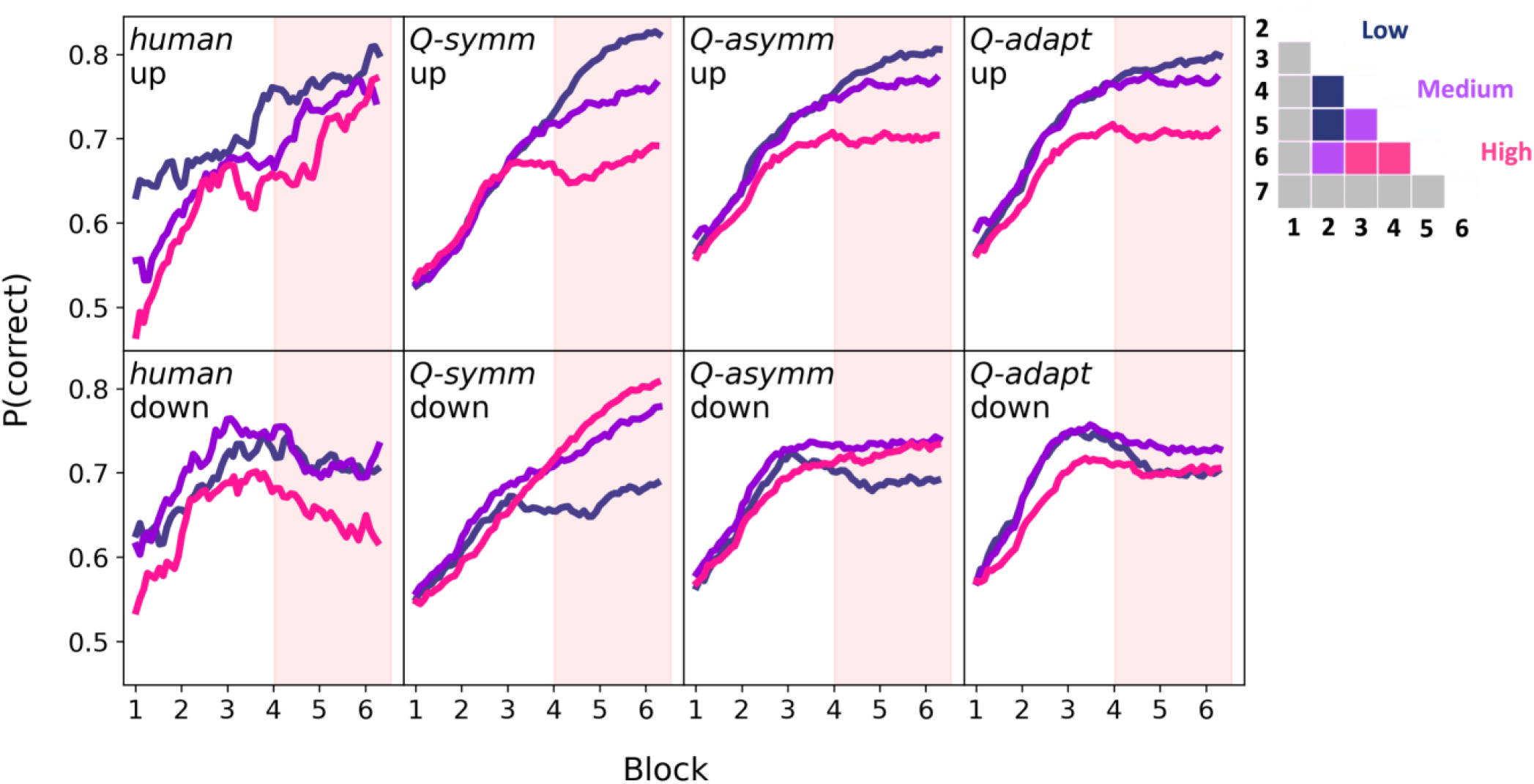
TI accuracy over the course of experiment in humans and fitted models. Mean accuracy for TI pairs was calculated using a sliding window of 100 trials. Red shaded half of each panel represents the post-changepoint phase of the experiment. Dark blue, purple and pink colours respectively refer to low, medium, and high-valued TI comparisons, excluding anchors (see choice matrix in legend). Humans (leftmost column) exhibited a differential impact of the changepoint on TI performance: whereas accuracy continued to improve in the ‘up’ group (upper leftmost panel), post-changepoint accuracy was disrupted in the ‘down’ group (lower leftmost panel). Simulating each candidate model using each participant’s best-fitting parameters revealed that whereas the asymmetric and adaptive models *Q-asymm* and *Q-adapt* (third and fourth columns, respectively) qualitatively reproduced this interaction effect, the symmetric model *Q-symm* performed equally well in both conditions.

We next examined the extent to which participants’ choice behaviour in the pre-changepoint period may have been reflective of a compressed latent value structure, a key signature of an asymmetric learning policy. Inspecting participants’ pairwise choice matrices (Fig 3A, left panels) showed evidence of value compression, such that lower-valued TI pairs (that is, pairs of items closer towards the top-left corner of the choice matrix) tended to be judged more accurately than higher-valued TI pairs (that is, pairs of items closer towards the bottom-right corner of the choice matrix). We quantified the slope of this compression effect using linear regression (Fig 4A and S2). Participants in both groups tended to exhibit asymmetry slopes significantly below 0, such that increases in combined pair value on TI trials were associated with a decline in accuracy (‘up’: mean *β* = -0.02 ± 0.01 SE, *t*(38) = -3.39, *p* < .001; ‘down’: mean *β* = -0.02 ± 0.01 SE, *t*(43) = -3.62, *p* < .001). This degree of asymmetry did not significantly differ between groups (*t*(81) = 0.10, *p* = .918). In line with previous work, we therefore found evidence that during the initial pre-changepoint phase, participants acquired a compressed value structure, consistent with an asymmetric learning strategy.

**Fig 3A-B.**
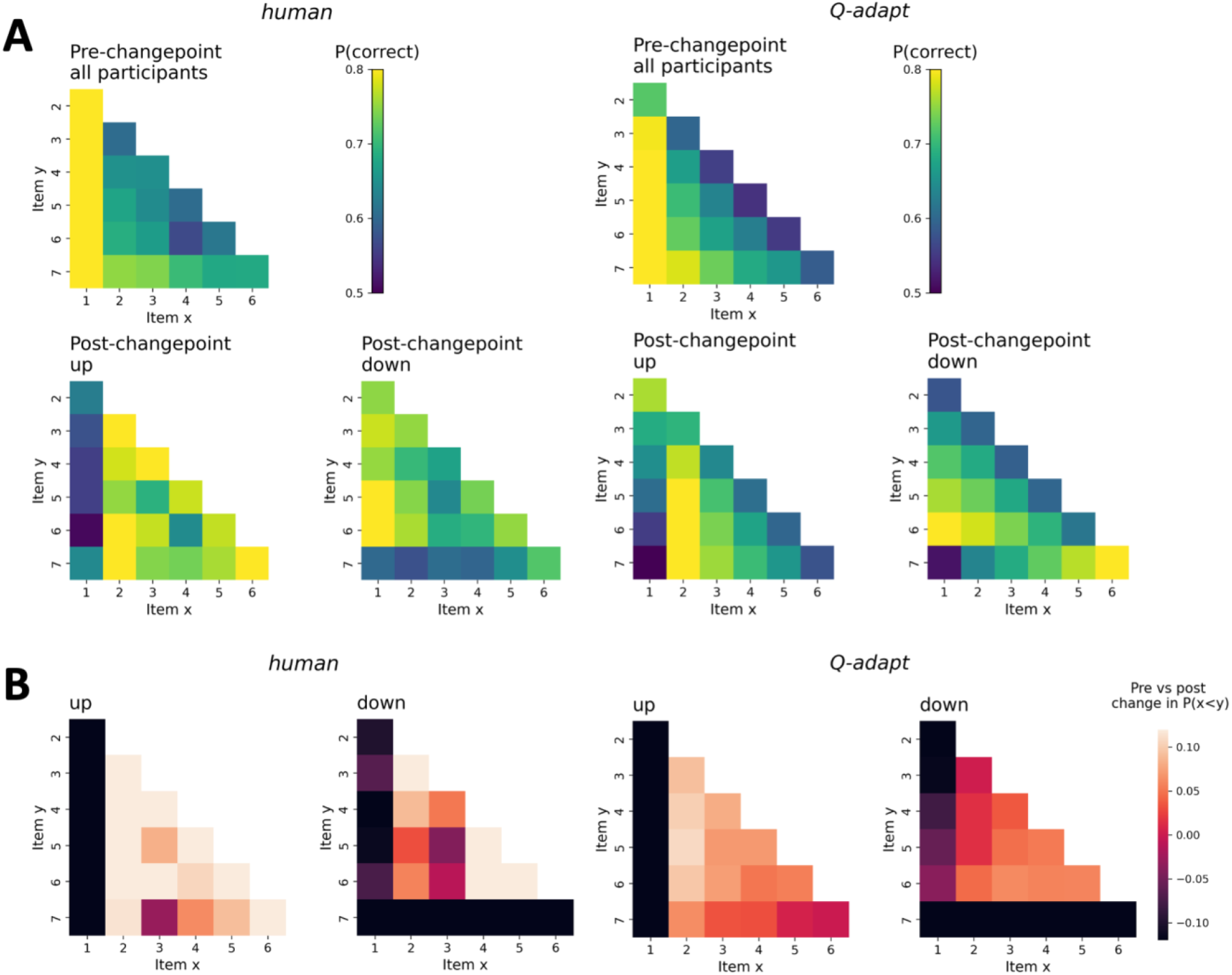
Choice matrices for humans (left panels) and the best-fitting model *Q-adapt* (right panels). **A)** Mean probability of choosing the *correct* item for each possible pairing, as represented by the colour-bar. Top row of panels displays pre-changepoint data collapsed across ‘up’ and ‘down’ participants, while the bottom row of panels splits post-changepoint data by group. **B)** Pre vs. post-changepoint change in P(x<y), i.e. the difference in preference for item *y* (matrix rows) over item *x* (matrix columns) from one changepoint to the next (note the change in metric compared to **A**). Lighter colours indicate that the agent’s preference for item y over x has increased, while darker colours indicate that it has decreased. Colour-bar value range was narrowed between -0.12 and 0.12 to improve legibility of differences among non-anchor pairs.

**Fig 4A-D.**
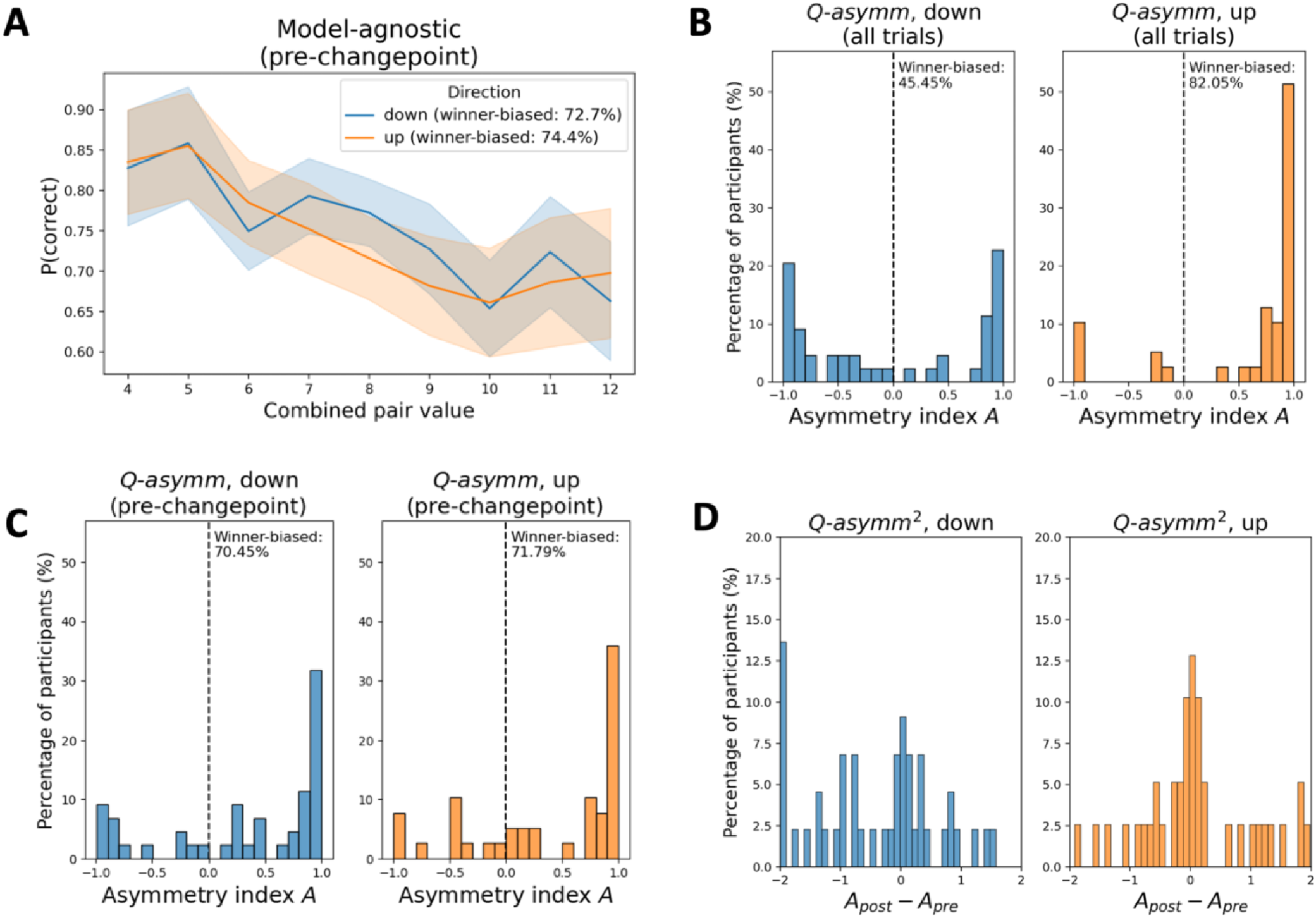
Model-agnostic and model-estimated learning asymmetry. **A**) The model-agnostic measure of participants’ learning asymmetry is the slope of the relationship between TI accuracy and combined pair value on all pre-changepoint trials (see also Fig S2). Participants with negative slopes are designated as winner-biased (see in-text legend for percentages). **B**) In contrast, the *Q-asymm*-based asymmetry measure refers to the normalised difference in best-fitting learning rates, where -1, 0 and +1 values for *A* indicate full loser bias, symmetry, and full winner bias respectively. Whereas ‘up’ participants tended to be strongly winner-biased when *Q-asymm* was fit to trials from the whole experiment (i.e. strong left-skew in right panel), ‘down’ participants were estimated to be more evenly split between winner- and loser-biased (i.e. bimodal distribution in left panel). The proportion of participants designated as winner- or loser-biased in the ‘down’ group according to this model-based metric therefore substantially deviated from that according to the model-agnostic metric in **A** (see in-plot percentages). **C**) In contrast, *Q-asymm* models fit to pre-changepoint trials were predominantly winner-biased in both groups. **D**) We fit *Q-asymm*^*2*^ to participant data, which was equivalent to *Q-asymm*, except that its two learning rates reset after the changepoint. Calculating the difference between the model’s pre- and post-changepoint asymmetry index *A* revealed a tendency to become less winner-biased in the ‘down’ group (left panel).

### Differential Impact of Changepoint on TI Performance

Turning to post-changepoint behaviour, we examined how effectively participants accommodated the different shifts in the ranking of one of the anchor items (i.e. *i*_*1*_ or *i*_*7*_) while preserving their knowledge about the remaining items (Fig 2, leftmost column). To isolate the impact of each changepoint on downstream inferential knowledge, and to avoid any skewing effect of pre-changepoint preferences for the moved anchor items, we focused on comparisons involving non-anchor items whose rank position had not changed in either group (i.e. from *i*_*2*_ to *i*_*6*_). Post-changepoint non-anchor accuracy was significantly above chance in both groups for adjacent pairs (‘up’: mean accuracy = 0.80 ± 0.02 SE, *t*(38) = 12.10, *p* < .001; ‘down’: mean accuracy = 0.73 ± 0.03 SE, *t*(43) = 8.48, *p* < .001), and for TI pairs (‘up’: mean accuracy = 0.74 ± 0.03 SE, *t*(38) = 7.03, *p* < .001; ‘down’: mean accuracy = 0.70 ± 0.03 SE, *t*(43) = 6.37, *p* < .001). To evaluate how accuracy developed from one phase of the experiment to the next, and whether these effects differed between groups, we conducted a series of 2 × 2 mixed ANOVAs with changepoint (pre vs. post) as a within-subjects factor, and direction (‘up’ vs. ‘down’) as a between-subjects factor. For adjacent pairs, we observed a significant main effect of changepoint (*F*(1,81) = 83.13, *p* < .001), reflecting a significant increase in accuracy from the first to the second half of the experiment (pre-changepoint: mean accuracy = 0.62 ± 0.01 SE; post-changepoint: mean accuracy = 0.76 ± 0.02 SE). Adjacent trial accuracy did not significantly differ between direction groups across the whole experiment (‘up’: mean accuracy = 0.72 ± 0.02 SE; ‘down’: mean accuracy = 0.67 ± 0.02 SE; *F*(1,81) = 3.87, *p* = .053), nor was there a significant changepoint x direction interaction effect (*F*(1,81) = 1.25, *p* = .268). Repeating this 2 × 2 ANOVA on TI accuracy, we likewise observed a main effect of changepoint (*F*(1,81) = 20.00, *p* < .001), which was similarly driven by an improvement in TI accuracy from the pre-to the post-changepoint phase of the experiment (pre-changepoint: mean accuracy = 0.65 ± 0.02 SE; post-changepoint: mean accuracy = 0.72 ± 0.02 SE). While the main effect of direction on TI trial accuracy was non-significant (‘up’: mean accuracy = 0.68 ± 0.02 SE; ‘down’: mean accuracy = 0.69 ± 0.03 SE; *F*(1,81) < 0.01, *p* = .950), we observed a significant changepoint x direction interaction (*F*(1,81) = 5.87, *p* = .018). Bonferroni-corrected post-hoc comparisons revealed that while participants in the ‘up’ group exhibited a significant improvement in TI accuracy from the pre-to the post-changepoint phases (pre-changepoint: mean accuracy = 0.63 ± 0.03 SE; post-changepoint: mean accuracy = 0.74 ± 0.03 SE; *t*(38) = 5.19, *p* < .001), participants in the ‘down’ group showed no such effect (pre-changepoint: mean accuracy = 0.67 ± 0.02 SE; post-changepoint: mean accuracy = 0.70 ± 0.03 SE; *t*(43) = 1.51, *p* = .277).

To inspect any differences in the development in TI accuracy after the changepoint more closely, we divided the post-changepoint phase in half and performed a further 2 × 2 ANOVA on non-anchor TI accuracy, but this time using these two halves of the post-changepoint data as the within-subjects factor, as opposed to pre-vs. post-changepoint. We observed no significant main effect of this factor (first half: mean accuracy = 0.71 ± 0.02 SE; second half: mean accuracy = 0.73 ± 0.02 SE; *F*(1,81) = 1.99, *p* = .162). However, the direction x post-changepoint half interaction effect was significant (*F*(1,81) = 6.39, *p* = .013). Bonferroni-corrected post-hoc comparisons revealed that this was similarly driven by a significant improvement in TI accuracy among the ‘up’ group from the first half of the post-changepoint phase to the next (first half: mean accuracy = 0.70 ± 0.04 SE; second half: mean accuracy = 0.77 ± 0.03 SE; *t*(38) = 3.03, *p* = .009), and a non-significant difference between the post-changepoint halves among ‘down’ participants (first half: mean accuracy = 0.71 ± 0.03 SE; second half: mean accuracy = 0.70 ± 0.04 SE; *t*(43) = 0.67, *p* > .999). Together, this indicates that the changepoint manipulation differentially impacted participants’ ability to infer transitive relations among unchanged items: while participants continued to improve non-anchor TI accuracy when *i*_*1*_ moved to the top of the hierarchy, non-anchor TI learning was relatively stunted in participants for whom *i*_*7*_ moved to the bottom of the hierarchy.

We next investigated the extent to which participants appropriately switched their choice preferences for whichever anchor item had moved to the other end of the hierarchy after the changepoint - i.e. P(choose *i*_*1*_) for ‘up’ participants, and P(choose *i*_*7*_) for ‘down’ participants (note: we excluded *i*_*1*_ vs. *i*_*7*_ trials from this analysis in order to isolate any changes in preference for these moved anchors with respect to the non-anchor items). In ‘up’ participants, we observed a significant increase in participants’ preference for the moved anchor *i*_*1*_ after the changepoint (pre-changepoint: mean = 0.15 ± 0.02 SE; post-changepoint: mean = 0.57 ± 0.06 SE; *t*(38) = 6.57, *p* < .001), and likewise a significant decrease in ‘down’ participants’ tendency to choose *i*_*7*_ after the changepoint (pre-changepoint: mean = 0.70 ± 0.03 SE; post-changepoint: mean = 0.38 ± 0.05 SE; *t*(43) = -7.49, *p* < .001). The *absolute difference* in choice preferences for the moved anchor before and after the changepoint did not significantly differ between the two groups (‘up’: mean difference 0.41 ± 0.06 SE; ‘down’: mean difference = 0.32 ± 0.04 SE; *t*(81) = 1.19, *p* = .238). Thus, both groups of participants appeared equally capable of correctly re-positioning whichever anchor item had moved to the other end of the hierarchy. This may suggest a certain degree of symmetry in updating the anchor items themselves after the changepoint, alongside the more general asymmetric updating of all other items, a possibility that we return to later in the *Results* section.

### Model Asymmetry

The foregoing behavioural analyses suggest that participants exhibited value compression effects and differential sensitivity to changes in relational structure consistent with a winner-biased belief-updating policy. Next, we fitted our symmetric and asymmetric RL models (*Q-symm* and *Q-asymm*) to the human experiment data, using the Akaike Information Criterion (AIC) to compare relative model fits (see *Model and Parameter Recovery* in *Materials and Methods* and Fig S3A-B) [29]. In accordance with Bayesian model selection approaches, we also calculated the protected exceedance probability (pxp) associated with each model, which quantifies the probability that a given model is the most frequent data-generating model of the entire set of candidates [30]. In both groups of participants, *Q-asymm* provided a better fit to participants’ behaviour than *Q-symm*, as confirmed using Wilcoxon signed-rank tests of AICs (‘down’: mean *Q-asymm* AIC = 349.81 ± 10.46 SE; mean *Q-symm* AIC = 366.12 ± 9.71 SE; *Z* = 5.22, *p* < .001; ‘up’: mean *Q-asymm* AIC = 339.26 ± 12.48 SE; mean *Q-symm* AIC = 363.53 ± 10.99 SE; *Z* = 5.25, *p* < .001) (Fig 5A). Comparison of pxps likewise revealed, in both groups of participants, a clear advantage for *Q-asymm* over *Q-symm* (‘up’: *Q-asymm* pxp > 0.99, *Q-symm* pxp < 0.01; ‘down’: *Q-asymm* pxp > 0.99, *Q-symm* pxp < 0.01). These initial model comparison analyses therefore not only replicate previously observed learning asymmetries, but also suggest that the differential impact of the changepoint in our modified TI setting is likewise best captured by the asymmetric learning agent *Q-asymm*.

**Fig 5A-B.**
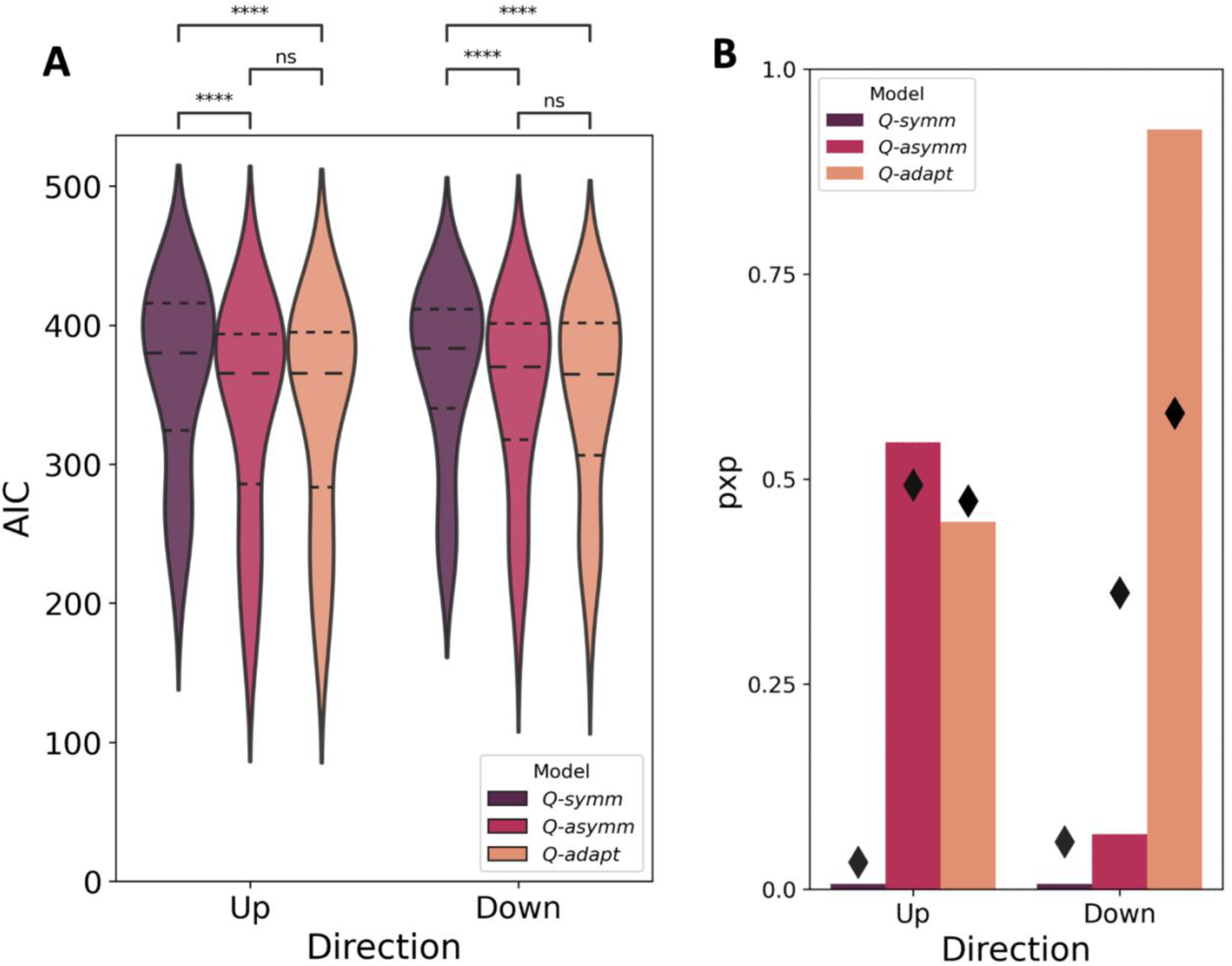
Model comparison for each candidate model, within each task condition. **A)** Lower AIC values indicate better fit of the model to the behavioural data. Dashed lines indicate quartiles of the data, while asterisks indicate a significant difference between AIC values for a given pair of models (i.e. *p* < .05; Wilcoxon signed-rank tests). **B)** Higher pxp values (bars) indicate greater probability that a given model is the most frequent data-generating model in the studied population, while diamonds indicate the estimated frequency of each model.

We next examined the model-estimated asymmetry index *A* of each participant under the *Q-asymm* model, where values of *A* closer to 1 or -1 indicate greater winner or loser biases respectively, and *A* = 0 indicates perfect symmetry between learning rates (see Eq 3. in *Materials and Methods*, ‘Behavioural Models’). As in previous work [5], values of *A* tended to be left-skewed in the ‘up’ group, indicating a strongly winner-biased learning asymmetry (Fig 4B, right panel). In addition to this majority of ‘up’ participants who were estimated to be winner-biased (N=32/39), there was also a small sub-group of participants for whom *A* was lower than 0, and hence who were estimated to be loser-biased under the best-fitting *Q-asymm* model (N=7/39). In contrast, *A* values for ‘down’ participants exhibited a more starkly bimodal distribution, such that participants were more evenly split between being either strongly winner-biased (N=20/44) or loser-biased (N=24/44) (Fig 4B, left panel). Indeed, non-parametric statistical comparisons revealed significantly lower values of *A* in ‘down’ participants compared to ‘up’ participants (‘down’: mean *A* = -0.02 ± 0.12 SE; ‘up’: mean *A* = 0.62 ± 0.10 SE; Mann-Whitney-U-test: *U* = 1269.00, *p* < .001). In contrast, when fitting *Q-asymm* to participants’ *pre-changepoint* choices only, we observed no significant difference in model-estimated asymmetry (‘down’: mean *A* = 0.34 ± 0.11 SE; ‘up’: mean *A* = 0.38 ± 0.11 SE; Mann-Whitney-U-test: *U* = 876.00, *p* = 0.873). The *A* values obtained from these pre-changepoint fits instead tended to be similarly left-skewed in both groups, providing estimates for the number of winner and loser-biased participants (**‘**up’: winner-biased N=28, loser-biased N=11; ‘down’: winner-biased N=31, loser-biased N=13) that more closely matched those obtained under our model-agnostic asymmetry slope metric reported earlier (cf. Fig 4A and S2). This suggests that while participants’ pre-changepoint behaviour may be best explained by a winner-biased learning policy, our model fitting procedure may have biased *Q-asymm*-derived learning rates towards capturing post-changepoint behaviour, leading to inflated estimates of loser learning rates. We address this possibility in the following section.

Our original hypothesis was that a differential impact of the changepoint on TI performance would arise as a direct consequence of the agent’s asymmetric learning policy – that is, the relative ease (or difficulty) in accommodating the ‘up’ (or ‘down’) relational change should be a function of each agent’s tendency to preferentially update winners or losers, up until the changepoint is reached. Such hypotheses were therefore derived under the assumption of a static degree of asymmetry, whereby each agent’s preferential updating of winners (or losers) remained constant over the course of the task, even in the face of the changepoint. However, it is also important to consider the possibility that such asymmetries may have varied over time as learning progressed, or as a function of objective changes in the task (namely, the changepoint). To evaluate the possibility that participants’ degree of learning asymmetry may have differed before and after the changepoint, we fitted a variant of *Q-asymm* equipped with two separate pairs of learning rates for winners and losers for each experimental phase, i.e. *a*^*+*^_*pre*_, *a*^*-*^_*pre*_ and *a*^*+*^_*post*_, *a*^*-*^_*post*_. We then calculated the asymmetry indices *A*_*pre*_ and *A*_*post*_ of this model *Q-asymm*^*2*^ using each of these pairs of fitted learning rates (Fig 4D). Participants in the ‘up’ group showed a winner-biased learning asymmetry in the pre-changepoint phase that did not significantly differ between changepoints (mean *A*_*pre*_ = 0.42 ± 0.11 SE; mean *A*_*post*_ = 0.51 ± 0.10 SE; Wilcoxon signed-rank test: *Z* = 0.35, *p* = .727). However, participants in the ‘down’ group underwent a significant reduction in their winner-biased learning asymmetry after the changepoint (mean *A*_*pre*_ = 0.37 ± 0.10 SE; mean *A*_*post*_ = -0.02 ± 0.12 SE; Wilcoxon signed-rank test: *Z* = 2.04, *p* = .041).

Interestingly, the ‘down’ participants for whom this change in learning asymmetry was most pronounced tended to be those who exhibited relatively high post-changepoint performance. For instance, participants’ difference between *A*_*post*_ and *A*_*pre*_ under *Q-asymm*^*2*^ was significantly negatively correlated with their post-changepoint non-anchor TI accuracy, and hence with their capacity to respond to the changepoint while minimising disruption to the unchanged transitive hierarchy (*r* = -0.73, *p* < .001; Fig S4A). Likewise, this reduction in learning asymmetry after the changepoint was positively correlated with participants’ pre-versus post-changepoint change in preference for the moved anchor *i*_*7*_, such that participants who correctly reduced their preference for *i*_*7*_ tended to show a greater reduction in their winner-bias after the changepoint (*r* = 0.38, *p* = .010; Fig S4B). In contrast, no such significant relationship held for ‘up’ participants, neither with respect to their post-changepoint non-anchor TI accuracy (*r* = 0.22, *p* = .181), nor their change in preference for the moved anchor *i*_*1*_ (*r* = 0.03, *p* = .846). These findings lend further support to the idea that although the changepoint experienced by ‘down’ participants disrupted TI learning at the group level, well-performing participants were nonetheless capable of leveraging an adaptive reduction in winner-biased asymmetry to respond more appropriately to the change in ground truth.

### Adaptive Asymmetry

The foregoing model comparison analyses indicate that while *Q-asymm* provides a good overall fit to both groups of participants’ behaviour, especially with respect to pre-changepoint trials, it is limited in its ability to account for well-performing participants who initially exhibited value compression, but who were nonetheless capable of responding appropriately to the downward change in relational structure. We therefore sought to explore how *Q-asymm* might be modified to make its learning policy flexible enough to capture the behaviour of such participants.

Inspiration for how differing degrees of asymmetry may arise as a function of some relevant task feature came from Ciranka et al.’s [5] finding that the sparsity of feedback appears to play a role in modulating learning policy asymmetry. Specifically, they observed that whereas participants tended to exhibit asymmetric belief-updating policies in the standard partial feedback TI paradigm, performance in a task offering full feedback on *all* comparisons, as opposed to just comparisons between neighbours, was best characterised by symmetric learning rates, and hence best fit by the symmetric model *Q-symm*. Such feedback regimes facilitate learning because they offer participants the opportunity to learn the cnarciness relations between non-neighbouring items directly. This also provides many more opportunities for the agent to confirm or revise their prior beliefs about the ordinal positions of the item set, which may lend itself to the application of symmetric updates to both compared items on a given trial. In contrast, in partial feedback settings where participants are required to ‘build’ a representation of the transitive hierarchy purely endogenously, the paucity of feedback that verifies or falsifies the agent’s beliefs about the ranking of items may necessitate asymmetrically prioritising the update of just one of the two compared items on a given comparison until a clearer representation of the item hierarchy has been formed.

We therefore formalised an adaptive agent *Q-adapt*, whose degree of asymmetry varied on a trial-by-trial basis as a function of the strength or uncertainty of the agent’s belief regarding the cnarciness relation between the two compared items. The rationale was that trials for which the agent’s belief about the two items is less certain may induce them to (asymmetrically) allocate a larger proportion of the overall update to one of the items. In contrast, on trials where the agent has a stronger belief, the receipt of feedback should provide a clear indication that this prior belief needs to be further reinforced or reversed via a more symmetrically distributed updating of both items. Drawing on the information theoretic notion of choice entropy, we derived an asymmetry variable *λ* which reflects the absolute strength of belief about the current comparison, and controls the degree to which the agent’s ‘base’ learning rate resource *a*^*0*^ is shared between *a*^*+*^ and *a*^*-*^ (Fig 6A; see Eqs. 6-8 in *Materials and Methods*, ‘Behavioural Models’). For example, assuming an agent with a general tendency towards winner-biased updates, when *λ* is 1 (indicating a weak preference), all of *a*^*0*^ will be allocated to *a*^*+*^, whereas *a*^*-*^ is set to 0. As *λ* approaches 0 (indicating a stronger preference), however, *a*^*0*^ is more evenly spread across both learning rates, meaning *a*^*+*^ and *a*^*-*^ become more symmetrical. Thus, whereas *Q-asymm* defines *a*^*+*^ and *a*^*-*^ as two free parameters, *Q-adapt* has a single base learning rate parameter *a*^*0*^ that is adaptively spread between *a*^*+*^ and *a*^*-*^ as a function of *λ* on a trial-by-trial basis.

**Fig 6A-C.**
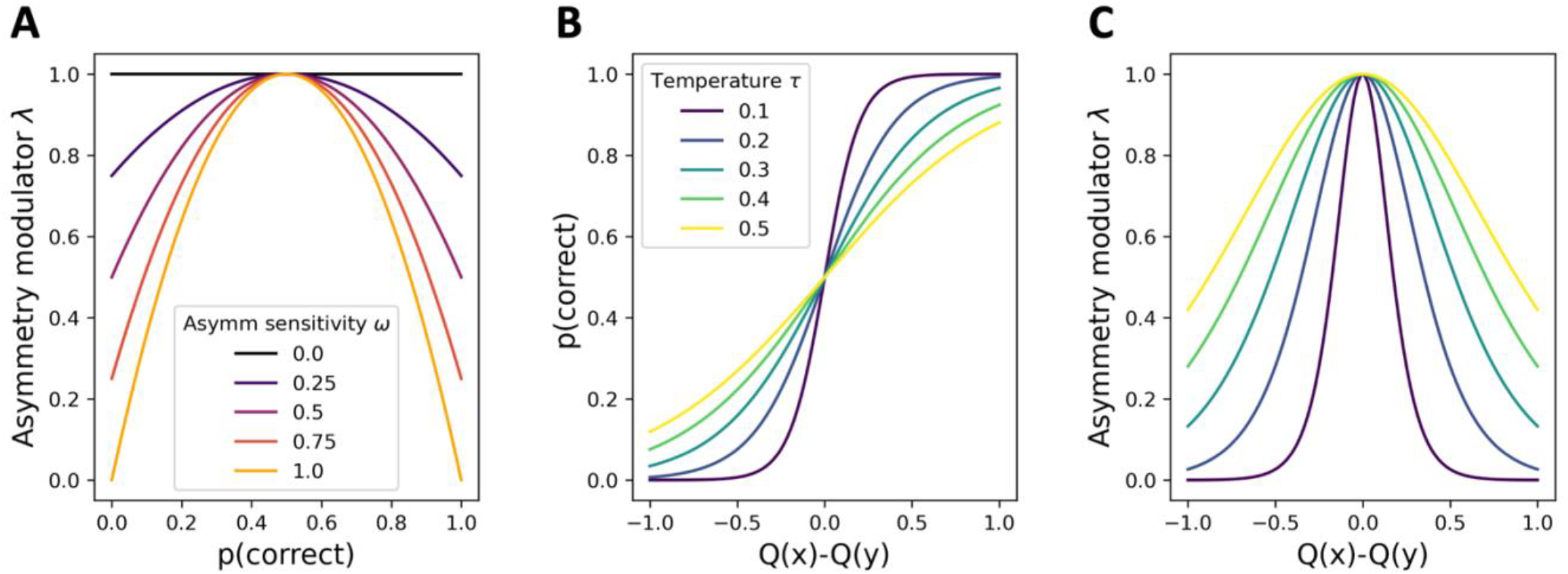
Illustration of how *Q-adapt* modulates its degree of learning asymmetry. **A)** The asymmetry modulator *λ* is given by a quadratic function of the agent’s preference strength – that is, the probability that they will choose *i*_*x*_>*i*_*y*_ on a given trial. The steepness of the asymmetry modulator function – that is, the degree to which *λ* is sensitive to changes in choice probability – is modulated by *ω*. **B)** Preference strength is a logistic choice function of the difference in value estimates for the compared items, the slope of which is determined by the temperature parameter *τ*. **C)** Assuming a constant *ω* (here, *ω*=1), then, given the relationship between *λ* and choice preference in **A**, which is itself dependent on *τ*, this means that the extent to which a difference in value estimates results in a smaller value of *λ*, and hence a more symmetric learning update, is at least partially shaped by each agent’s value for *τ*, and hence by their decision noise. In practice, *Q-adapt*’s learning dynamics can be roughly described as follows: at the beginning of the experiment, item values are not distinguishable, causing the agent to update items asymmetrically. As learning progresses and stronger preferences are formed, the agent begins to utilise a more symmetric update. Lower noise agents will exhibit a stronger tendency in this direction, meaning that, upon receipt of the *i*_*7*_<*i*_*1*_ feedback, they will more appropriately update these items (and indeed items on following trials) in a symmetric fashion, and thus resolve the ‘down’ changepoint with less difficulty. In contrast, higher noise agents will tend to update more asymmetrically across all value differences, leading to inflexible adaptation to the ‘down’ changepoint among those who are winner-biased.

In dynamically distributing learning updates in this way, *Q-adapt* models participants as tending to be more asymmetric in their updates towards the beginning of the experiment while they are still learning the transitive hierarchy, thus mirroring *Q-asymm’s* asymmetric policy. As learning progresses, and hence stronger (and, ideally, correct) beliefs about item relations are formed, learning updates are distributed more symmetrically (note that as the agent’s expectations about item relations become more accurate, this will in turn reduce the relative difference between predicted item values, resulting in a concomitant reduction in learning, as per Eq. 4). Once the changepoint is reached and the agent observes that *i*_*7*_<*i*_*1*_ – i.e. an outcome that contradicts the agent’s strong prior belief that *i*_*1*_<*i*_*7*_ –, the symmetric nature of the quadratic function allows for an updating of both *Q(i*_*1*_) and *Q(i*_*7*_) that is itself more symmetric, albeit still winner-biased. This is consistent with our finding that participants of both groups were equally capable of repositioning the moved anchor in each case, despite the differential impact of the changepoint on downstream TI performance.

The extent to which an agent may tend towards such symmetric updates is not only shaped by an additional sensitivity parameter *ω* (see Eq. 6), but also depends on how readily the agent forms strong preferences. This is itself determined by several interacting factors, including the rate at which the agent updates items upon receipt of new feedback (i.e. the learning rate), and the behavioural variability arising from the decision process (i.e. the temperature parameter *τ* of the logistic choice function; see Eq. 5 in *Materials and Methods*, ‘Behavioural Models’). In the present case, well-performing agents, such as those with lower values of *τ* will tend to more readily translate differences in value estimates into stronger choice preferences (Fig 6B), and hence will be more inclined to distribute more symmetric updates as learning progresses via lower values of *λ* (Fig 6C). In contrast, noisier agents will tend towards more asymmetric updates, limiting their ability to adapt to the change in relational structure occurring in the ‘down’ group. Thus, in modulating learning asymmetry as a function of choice preference, which is itself shaped by internal learning and noise parameters, the present implementation of *Q-adapt* allows agents to a) initially exhibit asymmetric learning while choice preferences are being acquired, and (crucially), b) appropriately deploy more symmetric learning later on in the learning phase under ‘well-performing’ learning and choice parameterisations.

We fitted this modified model *Q-adapt* to participants’ choices over the whole experiment, and repeated the Bayesian model selection steps to calculate model pxps, given the addition of this new candidate model (Fig 5A-B). Among ‘up’ participants, the adaptive model *Q-adapt* did not significantly differ from *Q-asymm* in terms of AIC (*Q-adapt*: mean AIC = 339.20 ± 12.65 SE; *Q-asymm*: mean AIC = 339.26 ± 12.48 SE; Wilcoxon signed-rank test of AICs: *Z* = 0.27, *p* = .791), and did not outperform *Q-asymm* in terms of pxp (*Q-adapt*: pxp = 0.45; *Q-asymm*: pxp = 0.54; *Q-symm*: pxp < 0.01). Among ‘down’ participants, *Q-adapt* yielded a slight but non-significant improvement in terms of AIC (*Q-adapt*: mean AIC = 348.37 ± 10.60 SE; *Q-asymm*: mean AIC = 349.81 ± 10.46 SE; Wilcoxon signed-rank test of AICs: *Z* = 0.81, *p* = .421), but clearly outperformed its static counterparts in terms of pxp (*Q-adapt*: pxp = 0.93; *Q-asymm*: pxp = 0.07; *Q-symm*: pxp < 0.01). Together, this indicates a narrow advantage for *Q-adapt* over *Q-asymm* in terms of model fit, particularly with respect to ‘down’ participants.

As a final model validation step, we simulated *Q-adapt* (along with all other models) using the best-fitting empirical parameters to verify whether this model was capable of qualitatively reproducing the key behavioural effects observed in our empirical dataset [29,31]. We first examined the consistency of the human and model-estimated value compression effects. In line with the descriptive results (cf. Fig 3A, upper left panel), *Q-adapt’s* pre-changepoint TI performance was characterised by a compressed value structure, with asymmetry slopes significantly below 0 (‘up’: mean *β* = -0.02 ± 0.01 SE, *t*(38) = -3.89, *p* < .001; ‘down’: mean *β* = -0.01 ± 0.01 SE, *t*(43) = -2.44, *p* = .019). Identifying participants as winner-or loser-biased according to the sign of their best-fitting *a*^*0*^ value, *Q-adapt* likewise yielded estimates for the proportion of participants falling into each category that were more closely in line with those gleaned from the sign of participants’ asymmetry slope (number of winner-biased participants under *Q-adapt:* ‘down’: 25/44 participants; ‘up’: 33/39 participants; cf. Fig 4B and S2). This stands in contrast to *Q-asymm*, which failed to reproduce a significantly negative asymmetry slope among ‘down’ participants (‘up’: mean *β* = -0.02 ± 0.01 SE, *t*(38) = -4.03, *p* < .001; ‘down’: mean *β* = -0.01 ± 0.01 SE, *t*(43) = -1.44, *p* = .157), while also underestimating the proportion of winner-biased participants according to its model-based asymmetry index *A*, as reported earlier.

Turning to model behaviour as a function of the changepoint, *Q-adapt*’s TI performance was differentially impacted by the change in underlying ground truth rankings, as in our behavioural data (cf. Fig 3B, left panels): non-anchor TI accuracy was relatively stunted in the ‘down’ group after the changepoint, whereas performance continued to improve in the ‘up’ group (Fig 3B, right panels). Interestingly, the exact pattern of TI disruption in ‘down’ participants deviated from that predicted by *Q-adapt* (and indeed *Q-asymm*); while our models predicted a more pronounced decline in lower-valued comparisons, the detrimental impact of the ‘down’ changepoint tended to be more strongly reflected in higher-valued comparisons (Fig 2, lower-leftmost and lower-rightmost panels). Nonetheless, as in our behavioural data, the broad pattern of a differential impact of the changepoint on inferential knowledge was supported by a significant changepoint x direction interaction effect on *Q-adapt*’s non-anchor TI accuracy (*F*(1,81) = 4.17, *p* = .044). This was driven by a significant improvement in TI accuracy from pre-to post-changepoint for ‘up’ participants modelled by *Q-adapt* (pre-changepoint: mean accuracy = 0.67 ± 0.02 SE; post-changepoint: mean accuracy = 0.75 ± 0.03 SE; *t*(38) = 6.56, *p* < .001), in contrast to a far less pronounced, albeit still significant, increase in TI accuracy between changepoints for ‘down’ participants modelled by *Q-adapt* (pre-changepoint: mean accuracy = 0.68 ± 0.02 SE; post-changepoint: mean accuracy = 0.72 ± 0.02 SE; *t*(43) = 2.43, *p* = 0.039).

Thus, considering not only our models’ predictive performance, as approximated by model evidence metrics, but also their ability to generate patterns of behaviour resembling those observed in humans, *Q-adapt* emerged as the model that best captured human TI performance.

## Discussion

TI is an instance of humans’ and other animals’ impressive ability to utilise knowledge gained about local relations to infer global, unseen relationships. By introducing different changes in relational structure, we demonstrated that winner-biased belief-updating confers different levels of flexibility to adapt to such changes in ground truth orderings: whereas relocating the worst item ‘up’ to the top of the hierarchy is readily accommodated, relocating the best item ‘down’ to the bottom has a more disruptive impact on downstream inferential knowledge.

Participants’ reduction in sensitivity to pre-changepoint TI comparisons with increasing combined value replicates compression effects previously observed in inferential learning settings [5]. Besides further underscoring the utility of using an RL framework to capture TI learning dynamics [4,5,32], we extend these findings by observing differences in adaptability to changes in relational structure that are consistent with an asymmetric, rather than symmetric, learning policy. Our findings lend further credence to the hypothesis that belief-updating asymmetries extend beyond two-armed bandit and foraging task contexts [10]. We note that the specific form of positivity bias in the present study is somewhat different to those investigated in the wider literature. In other RL paradigms, ‘positivity’ refers to the preferential update of values or options upon receipt of a positive (as opposed to negative) reward prediction error (RPE). Here, in contrast, the bias lies in the disproportionate updating of the winner and loser of a given binary comparison, independent of the sign of the RPE.

Our paradigm’s minimal change in underlying ground truth structure halfway through the task was reflected in a slight change in feedback that only subtly differed between groups: both sets of participants were given a single new piece of feedback (i.e. *i*_*7*_<*i*_*1*_), and only differed in the single comparison pair that no longer offered feedback (i.e. *i*_*1*_ vs. *i*_*2*_ for ‘up’ participants, and *i*_*6* vs._ *i*_*7*_ for ‘down’ participants). To model the updating of item value estimates in response to choice feedback, we assumed a relatively simple RL framework that only updated its cached value estimates for the currently presented pairs of items on receipt of feedback. Indeed, the utility of this ‘model-free’ approach in capturing human TI behaviour demonstrates that such inferential capabilities can proceed without necessarily invoking any abstract knowledge of the structural regularities entailed by particular relations (i.e. knowing that A<C *because* A<B and B<C). Nonetheless, it remains an intriguing possibility that humans could resolve the ambiguity initially induced by the changepoint by learning from trials from which they receive no feedback. In the present case, for example, a participant in the ‘up’ group might have learned to expect feedback, given the presentation of *i*_*1*_ *vs. i*_*2*_. The subsequent, unexpected omission of this feedback after the changepoint could induce them to update their value estimates for the presently compared items, and/or indeed items at the other end of the hierarchy, since it could be seen as diagnostic as to which of the underlying changes in ground truth explains the recently observed and highly surprising outcome *i*_*7*_<*i*_*1*_. This capacity to infer how the receipt or omission of feedback on a given comparison bears on items elsewhere in the hierarchy would therefore require a structural model of how the full set of items are related, potentially drawing on model-based approaches furnished with the ability to mentally simulate the outcomes of pairwise comparisons through replay [33–35].

The compression of participants’ learned value structures constitutes an instance of a more generalised distortion widely observed across psychophysical, numerical and economic decision-making contexts, whereby the discriminability between comparanda decreases with increasing stimulus intensity or magnitude [36–40]. Here, we propose that such compressed representations may emerge from an asymmetric learning policy (see also [5]). Nonetheless, we by no means argue that belief-updating biases are the only source of these ubiquitously observed psychometric distortions. Indeed, we note that the reduction in discriminability owing to increased overall value estimates across the hierarchy is not inconsistent with the view that compressed judgements of magnitude may arise, for example, from the mental organisation of numerical information on a power or logarithmic scale [38,39]. One potential way of disentangling the relative contributions of asymmetric policies and non-linear ‘Weber scaling’ of internal representations in the relational learning domain would be to more closely examine participants’ individual differences in the sign of the asymmetric learning bias: if the behavioural compression effects exhibited by winner-biased participants were equivalent to the anti-compression effects of loser-biased participants with equal *absolute* learning rate asymmetries, then this would further emphasise the role of asymmetric learning policies in the emergence of value compression. Relatedly, observing the opposite changepoint x direction interaction effect observed in our experiment, but among a predominantly *loser-biased* population – that is, disruption to inferential knowledge among the ‘up’ group, rather than the ‘down’ group – would lend further support to the idea that it is the sign of the learning asymmetry that is responsible for any (in-)efficient changepoint adaptation effects. Given the limited number of loser-biased participants in the present study, we leave this question for future work containing larger and more diverse samples of participants.

Our behavioural predictions were derived from *Q-asymm*, an RL agent that scaled its updates of winners and losers of pairwise comparisons according to asymmetric learning rates that remain fixed throughout the experiment. While this model significantly outperformed its symmetric counterpart *Q-symm*, it nonetheless underestimated the proportion of winner-biased participants. This raised the possibility that well-performing participants in the ‘down’ group were capable of both adapting to the change in relational structure, while also exhibiting pre-changepoint compression effects consistent with an initially winner-biased learning policy. We therefore introduced *Q-adapt*, a model whose trial-by-trial learning rate asymmetry varied as a function of the strength of its choice preference, thereby enabling well-performing participants to appropriately deploy more symmetric updating once the changepoint was reached. Existing models of changepoint adaptation typically possess the ability to separate the ‘aleatoric’ uncertainty pertaining to expected variability in an outcome from the ‘epistemic’ uncertainty arising from unexpected changes in a volatile reward environment [12–14]. Changepoints cause these models to increase their learning rates until the period of high epistemic uncertainty is resolved. *Q-adapt* lacks this capacity to track periods of volatility to modulate its overall learning rates, instead using the choice uncertainty on a given trial, as given by an entropy-like function, to directly modulate its degree of asymmetry.

*Q-adapt* parsimoniously unifies recent findings that asymmetric and symmetric learning policies each best explain human behaviour in, and indeed are optimal for, partial and full feedback TI regimes respectively [5]. We propose that the degree of belief-updating asymmetry flexibly varies according to the strength of an agent’s prior belief, and hence the informativeness, or entropy, of any resulting feedback. The formation of choice preferences from differences in value estimates is itself shaped by two model parameters: the amount of learning (as controlled by the base learning rate *a*^*0*^), and the decision noise with which learned item values are transformed into choices (as controlled by the temperature parameter *τ*; Fig 6B-C). The role of the latter parameter in asymmetry modulation dovetails with empirical work suggesting that magnitude compression effects and related psychometric distortions vary as a function of decision noise or task load [41–44]. Indeed, sensitivity to uncertainty has often been incorporated into RL frameworks in various guises, and has been suggested as guiding the use of different behavioural controllers in humans [45], the flexible combination of reward information in primates [46], and the volatility-induced adaptation of learning rates via meta-learning in rodents [47]. In the present case, the concept of uncertainty may more appropriately pertain to the agent’s prior confidence about the relative difference between item values at the time of choice. For example, if an agent has a stronger preference for *I*_*x*_<*I*_*y*_, and thus a less noisy representation of the relative values of these items, then the receipt of feedback that either confirms or disconfirms this belief may more unambiguously be incorporated into the agent’s value estimates in the form of a more symmetric update. In contrast, uncertain beliefs about less discriminable items may be associated with greater noise or working memory load, making it more appropriate to focus one’s update on just one item. Although our exploratory model comparison was intended to formalise the idea that stronger preferences should induce more symmetrical updates, there are several other task-related or internal variables that may covary with the strength of an agent’s preference, including confidence or surprise, the expected value of the chosen or unchosen option, the RPE magnitude, or the balance of exploration versus exploitation etc.. Future work could disentangle these candidate task features or decision-making variables that may give rise to fluctuating levels of asymmetry.

Theoretical accounts have proposed that the degree of learning rate asymmetry is optimally adapted to the richness of a reward environment, such that positive learning rate asymmetries maximise rewards in ‘poor’ environments, while negative asymmetries maximise rewards in ‘rich’ environments [11,48]. Asymmetric updating in response to relational feedback can be thought of as optimal in a similar way; under sparse feedback, prioritising the update of just one of the two compared items magnifies relative differences among item estimates during initial learning, and is therefore optimal for the ‘building’ of a value structure in which value estimates are clearly separated [5]. A corollary of these theoretical frameworks is that learning biases should dynamically *invert* as a function of the amount of reward available, although empirical evidence for such inversion is mixed [23,24; although see 25]. Our adaptive framework explores the possibility that the relative balance of positive and negative learning rates may dynamically *narrow* over time, rather than reverse. In any case, the mixed empirical evidence for adaptive asymmetries may be due to different operationalisations of reward richness. For instance, the above studies manipulated the average reward rate by controlling the probability that a reward would be received upon selection of one of two bandits. In contrast, the proportion of comparisons offering binary choice feedback, relative to those offering no feedback, did not change over the course of our TI changepoint task, nor did it vary between groups. Thus, in the present study, it is the trial-by-trial variability in choice preference strength, rather than the distribution of rewards, that is hypothesised to have an impact on learning rate asymmetry adaptation.

Aside from the RL framework deployed here, one can alternatively examine the TI changepoint problem under a Bayesian inference scheme, as has widely been done in the context of TI [3,32,49], and indeed structure learning more generally [50–52]. Under this broad class of frameworks, our TI changepoint scenario could be viewed as a problem of resolving feedback ambiguity: the new observation that *i*_*7*_<*i*_*1*_ is, at first, equally consistent with a change in *i*_*1*_’s ranking as it is with a change in *i*_*7*_’s ranking, meaning the agent must track the likelihood of subsequent choice feedback under each of these hypotheses about the new underlying ground truth structure. A wealth of literature has likewise researched the role of episodic processes, likely implemented in the hippocampus, in enabling inference and generalisation [53,54]. More specifically, TI may be supported by ‘retrieval-based’ inference mechanisms that reactivate and recombine pattern-separated representations of specific relations [35,55], or via a more ‘encoding-based’ recruitment of inferred relationships via overlapping structural representations [56,57]. Since the present study focused on how the reorganisation of relational knowledge intersects with widely observed biases in value learning, we did not incorporate into our models the possibility that transitive learning might also involve episodic memory processes [5,35]. Elucidating whether and how such episodic processes ‘feed into’ the caching of item values would therefore be a promising avenue for future work.

Several lines of research have connected elementary belief-updating biases with research in clinical settings. While positivity biases may provide an adaptive means of promoting positive well-being [58] or motivation [6], converging empirical and theoretical work has also implicated more pessimistic learning rates in a range symptoms of Major Depressive Disorder [59–61]. Our finding that belief-updating biases confer different levels of flexibility to changes in relational structure raises interesting questions about whether or not such differences in adaptability also cut across clinical populations. It would be particularly interesting to consider such asymmetries in changepoint adaptability in the context of risk-seeking or gambling behaviour, since they predict differences in the influence of various outcomes – e.g. a change in a previously low-performing bet versus a change in a previously high-performing bet – on reward expectations pertaining to unchanged bets. While our paradigm contained fully deterministic relational feedback, and therefore did not incorporate any risk or outcome variance per se, evidence suggests that the degree of learning asymmetry shapes the relationship between the environment’s reward variability and an individual’s tendency to seek or avoid risks [62–64]. Given our hypothesis that asymmetry dynamically varies as a function of preference strength, which in turn is influenced by an agent’s decision noise, it would be worthwhile to consider the role of belief-updating biases in value compression and changepoint adaptability effects under different levels of outcome variance (i.e. via probabilistic relational feedback), and how this relationship might be moderated by clinically relevant symptoms or traits.

Our RL agents constituted descriptive models of how biased learning policies give rise to subjective value distortions and differences in behavioural adaptability. While we make no causal or mechanistic claims about the dynamics of relational learning in the brain, research centring on neuromodulatory activity in the basal ganglia and brainstem may offer plausible accounts for how updates may be asymmetrically scaled during RL. Subpopulations of striatal neurons with distinct excitatory and inhibitory properties (i.e. D1 and D2 receptors, respectively) may provide a means of differential engagement of dopamine-mediated learning as a function of positive or negative prediction errors [65–67]. Likewise, empirical work has implicated serotonergic systems operating over behaviourally relevant timescales in the ability to track and adapt to changes in the volatility of reward environments [47,68]. It would therefore be interesting to consider how such neural accounts extend beyond bandit tasks to structure learning settings that more explicitly engage generalisation and inference processes. In addition, our investigation of differences in adaptability to changes in underlying relational structure ties into research exploring how neural and artificial systems reconfigure knowledge at the representational level in response to new information. Evidence suggests that the linking together of transitive hierarchies is mirrored in the joining of neural manifolds in fronto-parietal regions and deep neural networks alike [18]. Examining how the differences in relational adaptability observed in the present study might be recapitulated in a neural network may in turn yield neuroscientific hypotheses about how representational geometries may be (in-)efficiently reorganised in response to changes in environmental structure.

## Materials and Methods

### Participants

Participants (N=150) aged between 18-40 years were recruited online via Prolific Academic (74 female; mean age 27 ± 5.27 years SE). After confirming their written informed consent, participants were randomly allocated to one of two groups: the ‘up’ group (N=76; 37 female; mean age = 27.14 ± 5.12 years SE), or the ‘down’ group (N=74; 37 female; mean age = 26.85 ± 5.41 years SE). Participants received compensation of £6.00, plus a performance-dependent bonus of £2. The study was approved by the Ethics Committee of the Max Planck Institute for Human Development.

Since our study focused on the impact of the changepoint manipulation on learned knowledge, we implemented a performance-related inclusion criterion. Participants in both groups experienced the same trial structure before the changepoint was reached (albeit with different item allocations and trial sequences). We therefore used a binomial test to compute a performance threshold above which the likelihood that participants were performing at chance on *pre-changepoint trials* was 0.01 (i.e. following the criteria used by Ciranka et al. [5]), thus avoiding a confound by the experimental manipulation of interest. One additional participant was excluded for exhibiting a high proportion of missed responses (>60% of 322 trials). After the application of these criteria, N=83 participants (36 female; mean age = 26.90 ± 5.34 years SE) remained for analysis (‘up’: N=39; ‘down’: N=44). Restricting the application of this criterion to the first half of the experiment while participants were still learning to perform the task amounted to a somewhat conservative approach, in turn resulting in a relatively high proportion of participants being excluded. Nonetheless, we note that when we applied a more liberal threshold for inclusion (*a* = 0.1), which left N=103 participants (‘up’: N=53; ‘down’: N=50), our core findings – i.e. a differential impact of the changepoint on downstream TI performance, best explained by our adaptive asymmetry model *Q-adapt –* remained unchanged.

### Stimuli, Task and Procedure

The behavioural task was an adapted version of the TI paradigm used in Experiment 4 of Ciranka et al.’s study [5], and was programmed in PsychoPy 2022.2.2 [69]. Seven images of everyday objects and animals drawn from the BOSS database [70] were randomly assigned a ground truth rank from 1-7 at the beginning of the experiment for each participant. Participants were told that their task was to learn about how the items related to one another with respect to how ‘cnarcy’ they are. They were informed that whether or not an item was more or less cnarcy than another was unrelated to any characteristics that these items have in real life. Rather, participants could only learn about cnarciness through the feedback provided on each trial of the experiment.

On each trial, following a 0.5s fixation cross, two items were simultaneously presented on the left and right side of the screen on a white background for up to 2.5s. Participants were instructed to select whichever item they thought was more cnarcy than the other as accurately and as quickly as possible using the left or right arrow key. They were informed that, on some trials, they would receive on-screen feedback (“correct”/”incorrect”) about whether or not they had correctly chosen the cnarcier item. Unbeknownst to participants, the delivery of feedback was determined by the relative positions of the two items in the underlying cnarciness hierarchy that was established at the start of the experiment: if the two items were neighbouring in their rank (‘adjacent trials’, e.g. *i*_*3*_ *vs. i*_*4*_), then participants received feedback (“correct” or “incorrect”) about their choice for 0.5s, whereas if the items were non-neighbours (‘TI trials’, e.g. *i*_*3*_ *vs. i*_*5*_), then no feedback was provided. If no selection was made within 2.5s, a ‘missed response’ was recorded. Trials were separated by an inter-trial interval of 0.6s.

Using a set of seven items resulted in 21 possible stimulus pairings. Within each block, the six adjacent pairs were repeated four times, while TI pairs were repeated twice. This gave rise to a total of 54 trials per block, of which 24 provided feedback and 30 provided no feedback. The serial order of trials was pseudo-randomised, with left and right positions counterbalanced within each block.

The entire experiment consisted of six blocks, each followed by a short attention check. At the start of the experiment and before each block, participants were reminded that not all items would stay as cnarcy for the entire experiment. Rather, on some blocks, certain items may or (may not) become more or less cnarcy, meaning their relations to other items (as reflected in choice feedback) may change. In reality, such a change was only introduced in the fourth block, such that from this block onwards, the ground truth item hierarchy changed in a manner determined by the group to which participants had been assigned. In the ‘up’ group, the hitherto lowest-ranking item *i*_*1*_ moved ‘up’ the hierarchy to become the highest-ranking item, whereas in the ‘down’ group, the hitherto highest-ranking item *i*_*7*_ moved ‘down’ the hierarchy to become the lowest-ranking item. Given that choice feedback continued to only be delivered on trials comparing items of neighbouring rank, such changes in ground truth structure resulted in the following minimal changes to the feedback received by each group: 1) participants in both groups now received feedback informing them that *i*_*7*_<*i*_*1*_, 2) participants in the ‘up’ group now no longer received feedback that *i*_*1*_<*i*_*2*_, and 3) participants in the ‘down’ group now no longer received feedback that *i*_*6*_<*i*_*7*_. Thus, the changepoint meant that both groups learned about a single new relation, and only differed in the relation for which choice feedback was retained or withdrawn.

After the final block, participants performed three short tasks to test their explicit knowledge of the item hierarchy. First, using the mouse to drag and drop items, participants were asked to arrange the items according to how cnarcy they thought they were by the end of the experiment. Next, they were asked to click on whichever items (if any) they believed had changed how cnarcy they were at any point in the experiment. Finally, participants were given the opportunity to enter, using the keyboard, any comments they wanted to share on, for example, how they performed the task, the nature of the feedback received, or how difficult they thought the task was.

While we do not analyse this post-task data here, it can nonetheless be freely accessed alongside the rest of behavioural data (see Data Availability).

### Behavioural Models

We assume a simple Rescorla-Wagner learning rule to model how agents update their value estimates of items in response to relational feedback:

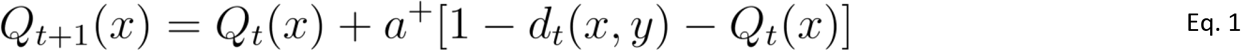

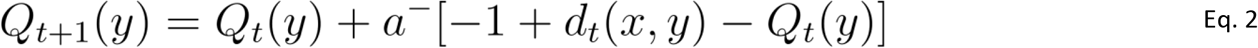

where *Q*_*t*_ is the estimated item value at time *t*, and *a*^*+*^ and *a*^*-*^ are the learning rates for the winning and losing items *x* and *y* respectively. For the symmetric agent *Q-symm, a*^*+*^ = *a*^*-*^, such that these learning rates are modelled as a single free parameter. In contrast, for the asymmetric agent *Q-asymm, a*^*+*^ and *a*^*-*^ can freely vary. This allowed us to obtain each participant’s model-estimated asymmetry index *A*, calculated as the normalised difference between best-fitting learning rates under *Q-asymm*:

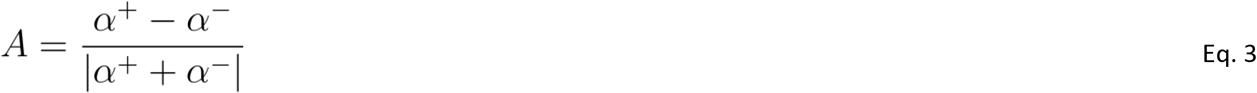

In the learning equations 1-2, *d*_*t*_*(x,y)* represents the relative difference between *Q*_*t*_*(x)* and *Q*_*t*_*(y)*, i.e.:

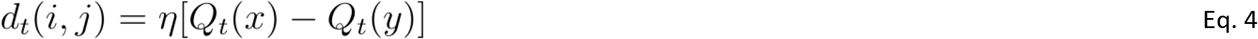

where *η* is a scaling factor. This formalises the assumption that value updates scale with the difference between estimated item values. For instance, if an agent observes that *i*_*x*_>*i*_*y*_, this outcome should only induce a small change in value estimates for these items if the agent had already learned to expect this outcome (i.e. if *Q(x) >> Q(y)*). In contrast, observing that *i*_*x*_<*i*_*y*_ would be highly surprising, given the agent’s existing beliefs about the relative values of these items, thus demanding a stronger update in the relevant value estimates. Incorporating such relational difference-weighting of value updates is necessary for *Q-symm* and *Q-asymm* to accomplish TI for non-anchor items (i.e. those of intermediate rank) [5]. We note that, depending on the value of the scaling factor *η*, the inclusion of the relative difference term *d*_*t*_ can ‘overflow’ the bounds (i.e. 1 and -1) of the Rescorla-Wagner rule in Eqs. 1-2 – that is, the term that is added to *Q(x)* in Eq. 1 or subtracted from *Q(y)* in Eq 2. may end up being negative or positive, respectively. In such cases, the estimate of the winner would therefore *decrease*, and/or the estimate of the loser would *increase*. In order to prevent such edge cases, we therefore apply a positive rectifier function to the winner update and a negative rectifier function to the loser update, such that any negative winner updates or positive loser updates are clipped at 0.

Finally, we used a logistic choice function to define the probability of choosing *i*_*x*_>*i*_*y*_ based on the difference between estimated item values:

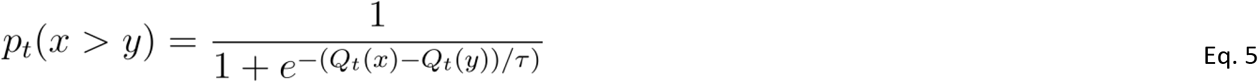

where *τ* is the temperature parameter determining the shape of the sigmoid function, and thus the degree of noise in choices based on item value differences.

The learning rates *a*^*+*^ and *a*^*-*^ remain static for *Q-symm* and *Q-asymm*. In contrast, the adaptive asymmetry *Q-adapt* is capable of modulating the degree to which learning updates are shared between a given comparison’s winner and loser on a trial-by-trial basis. On adjacent trials, we calculate an asymmetry modulator *λ*, bound between 0 and 1, as a quadratic function of the strength of the agent’s prior belief about how items *x* and *y* are related upon receipt of choice feedback:

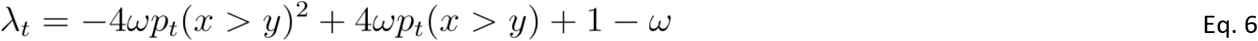

The value of *λ* is minimal, causing more symmetric updating, when an agent’s prior belief is strong and thus clearly supported or contradicted by the receipt of binary feedback (i.e. when *p(x<y)* approaches 1 or 0), whereas it is maximal, causing more asymmetric updating, when the agent has no preference (i.e. when *p(x>y)* = 0.5). *ω* is an additional sensitivity parameter bound between 0 and 1 controlling the shape of the quadratic asymmetry modulator function (Fig 6A). This determines how readily an agent adapts their degree of learning rate asymmetry as a function of the strength of their choice preference, effectively implementing a quadratic function that can be shallower or steeper depending on the value of *ω*. When *ω* is 0, the agent’s asymmetry is insensitive to changes in belief strength, such that *λ*=1 (i.e. full asymmetric updating) for all choice probabilities. When *ω* is 1, the Eq. 6 becomes roughly equivalent to a choice entropy function (cf. Eq. 9). (Note: best-fitting values for *ω* were bimodally distributed around 0 and 1 (i.e. corresponding to no adaptability and maximal adaptability of learning asymmetry, respectively), and did not significantly differ between groups (‘up’: mean *ω* = 0.37 ± 0.06 SE; ‘down’: mean *ω* = 0.47 ± 0.06 SE; Mann-Whitney U-test: *U* = 664.0, *p* = .077). This indicates that participants in both groups could be broadly divided into those whose learning asymmetry was or was not sensitive to changes in choice preference strength).

The *λ* term can then be used to distribute the agent’s base learning rate *a*^*0*^ across 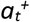 and 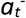 according to the following linear equations:

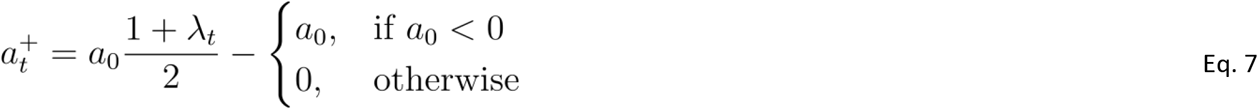

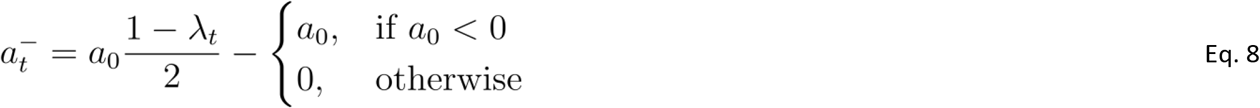

Since we allowed *a*^*0*^ to take on negative values, the inclusion of the rightmost term in Eqs. 7 and 8 enabled agents to vary in terms of whether their distribution of *a*^*0*^ across *a*^*+*^ and *a*^*-*^ was winner-biased (i.e. *a*^*0*^ > 0, and hence) or loser-biased (i.e. *a*^*0*^ < 0, and hence 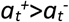). Note that we assume that agents cannot *reverse* their bias for the winners or losers of comparisons – for instance, for a winner-biased participant fit with *a*^*0*^ > 0, 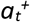 can only be greater or equal to 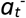. This is consistent with recent empirical work finding no evidence for an adaptive reversal of the sign of humans’ learning asymmetries [23,24; but see 25].

We also considered an alternative version to *Q-adapt* in which the asymmetry modulator *λ* is simply an entropy function of the choice preference strength, such that Eq. 6 is replaced by the following:

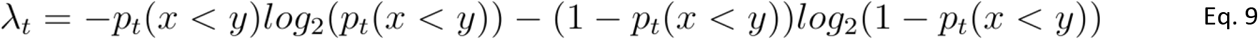

However, fitting this model to participant data revealed a significantly worse fit relative to the original ‘quadratic’ variant of *Q-adapt* described in Eq. 6 (‘quadratic’ *Q-adapt*: mean AIC = 344.06 ± 8.14 SE; ‘entropy’ *Q-adapt*: mean AIC = 348.11 ± 7.71 SE; Wilcoxon signed-rank test of AICs: *Z* = 2.29, *p* = .022). Given this inferior predictive performance for the ‘entropy’ model variant of *Q-adapt*, we excluded it from our formal model comparison.

### Model Fitting and Comparison

We estimated model parameters by minimising the log-likelihood of each model, given each participant’s single-trial responses. We used Scipy’s differential evolution method [71,72] over 500 iterations with the following lower and upper parameter bounds: *a*^*+*^/*a*^*-*^: (0;0.5); *a*^*0*^: (−0.5;0.5); *η*: (0;10); *τ*: (0;1). From the resulting log-likelihood values under these best-fitting parameter estimates, we computed AIC values as an approximation of model evidence, where lower AIC indicates better goodness of fit, while penalising for model complexity [73]:

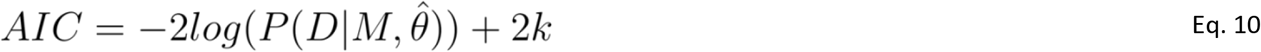

This amounts to the likelihood of a participant’s choice data *D* over the trials of interest, given a particular model *M* and its best-fitting parameters 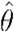, plus a penalty term *k* corresponding to the number of free parameters. AIC values were in turn used to quantify the protected exceedance probability (pxp) associated with each competing model using the Variational Bayesian Analysis toolbox in MATLAB [74]. pxp values amount to the probability that a given model fits participants’ data better than all other competing models, and hence is the most frequent data-generating model in the studied population. In contrast to the exceedance probability metric, the pxp additionally accounts for the null hypothesis that there is no difference in the frequencies of each model type [30].

### Model and Parameter Recovery

An important prerequisite for comparing models fitted to empirical data is that they are identifiable – that is, such models should behave in ways that renders them distinguishable under the selected model evidence metric [29]. To validate our model comparison approach, we first took each model’s best-fitting parameters for each of the 83 participants, and used these to generate 10 experimental runs of synthetic (binomial) choice data on each participant’s set of trial sequences. We then fitted each model to the generated data and evaluated how often each model provided the best fit. The resulting confusion matrix thus provides a measure of the conditional probability that a model fits the data best, given the true generative model: *p(fit*|*gen)*. In turn, this allows one to ‘invert’ the confusion matrix according to Bayes rule, under the assumption of a uniform prior over all models:

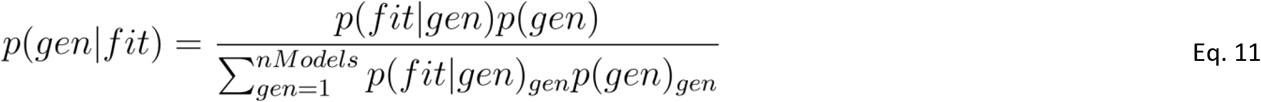

In quantifying the probability that the data were generated by a specific model, given that this model provided the best fit to the generated data, the inverted confusion matrix helpfully complements the model recovery procedure. As can be observed in the AIC-based confusion and inverted confusion matrices (Fig S3A-B), our models of interest exhibited robust recoverability in both ‘down’ and ‘up’ task settings. We note that model separability was greatly improved when using the AIC relative to when using (inverted) confusion matrices that used the Bayesian Information Criterion (BIC) [75], which over-penalised *Q-asymm’s* additional free parameter. Thus, we used AIC as our approximation of model evidence.

Finally, to validate inferences about empirical parameters obtained from our model fitting procedure, we simulated choice behaviour under our two key ‘winning’ models of interest, *Q-asymm* and *Q-adapt*, using the best-fitting parameter settings estimated for each participant, and then re-fit these models to these generated datasets [29]. We then repeated the process while incrementally varying each of the data-generating parameters over 10 evenly spaced values within the lower and upper bounds used for our model fitting procedure (see ‘Model Fitting and Comparison’). In both task conditions and model types, the ‘true’, data-generating parameters strongly correlated with their recovered counterparts (min *r* = 0.50, max *r* = 0.90), and only weakly correlated with all other recovered parameter types (min *r* = -0.29, max *r* = 0.13; Fig S5A-B). Likewise, we observed only weak correlations among the recovered parameters themselves (min *r* = -0.22, max *r* = 0.07), indicating that our fitting procedure did not introduce any ‘trading off’ among parameters, and thus further validating inferences drawn about these parameters (Fig S4C) [29,76].

## Author Contributions

The following list of author contributions is based on the CRediT taxonomy:

**Conceptualisation:** T.A.G., B.S.

**Data Curation:** T.A.G.

**Formal Analysis:** T.A.G.

**Funding Acquisition:** B.S.

**Investigation:** T.A.G.

**Methodology:** T.A.G., B.S.

**Project Administration:** T.A.G., B.S.

**Resources:** B.S.

**Software:** T.A.G.

**Supervision:** B.S.

**Validation:** T.A.G.

**Visualisation:** T.A.G.

**Writing - Original Draft:** T.A.G.

**Writing - Review & Editing:** T.A.G., B.S.

## Acknowledgements

We are grateful to Peter Dayan for valuable discussions and for comments on the manuscript. We also thank Christian Doeller for helpful feedback during project conceptualisation, and Philip Jakob and Maik Messerschmidt for technical support.

## Data Availability

The data that support the findings of this study are available at dx.doi.org/10.6084/m9.figshare.26147470

## Code Availability

The experiment and analysis code is available on GitHub at https://github.com/tgham/asymm_switch

## Supporting Information

**Fig S1. Predicted TI learning curves under simulations of symmetric and asymmetric RL models**. Same as Fig 2, but simulated using empirical parameter estimates for *Q-symm* (upper panels) and *Q-asymm* (lower panels) matching those observed by Ciranka et al. [5].

**Fig S2. Value compression effects in human behaviour**. Our model-agnostic measure of learning asymmetry – i.e. the slope of the relationship between combined item value and accuracy on pre-changepoint TI trials – was significantly lower than 0, indicating value compression. In-text percentages refer to the percentage of participants in each group whose asymmetry slope was below 0, thereby indicating those participants designated as winner-biased (cf. Fig 4A).

**Fig S3A-B. Model recovery analysis**. Our three candidate models generally exhibited good identifiability both in terms of *p(fit*|*gen)* (**A**) and *p(gen*|*fit)* (**B**). See *Materials and Methods, ‘Model and Parameter Recovery’* for details.

**Fig S4A-B. Changepoint-induced changes in learning asymmetry are related to behavioural performance**. Among ‘down’ participants, pre-vs. post-changepoint differences in learning asymmetry estimated with *Q-asymm*^*2*^ were significantly negatively correlated with post-changepoint non-anchor TI accuracy (**A**), and positively correlated their tendency to correctly change their preference for the moved anchor item (i.e. *i*_*7*_; **B**). In other words, participants in the ‘down’ group who appropriately adapted to the change in relational structure exhibited a larger reduction in their winner-biased learning asymmetry. In contrast, these relationships were non-significant among ‘up’ participants.

**Fig S5A-B. Parameter recovery analysis**. Our two winning models *Q-asymm* and *Q-adapt* exhibited strong parameter recovery. In other words, in the parameter correlation matrix in **B**, simulated parameters (columns) correlated strongly with their recovered counterparts (rows), resulting in strong correlation values along the diagonal. In contrast, simulated parameters correlated weakly with each other recovered parameter type (i.e. off-diagonal). In addition, we observed weak correlations *among* recovered parameters, as shown in **C**. See *Model and Parameter Recovery* in *Materials and Methods* for details.

