## Supplementary figures and images for "Asymmetric learning and adaptability to changes in relational structure during transitive inference"

### Fig_S1.png

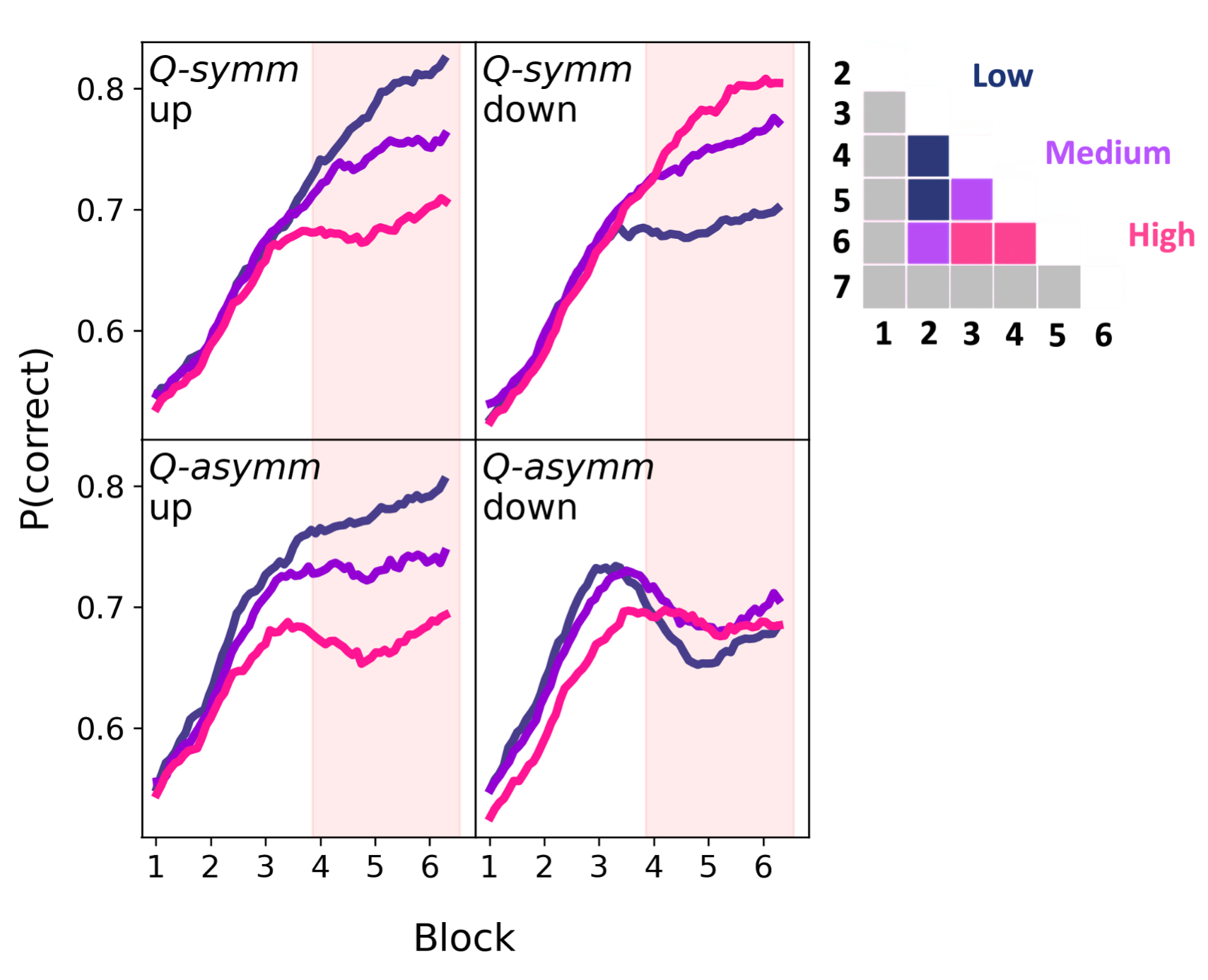

### Fig_S2.png

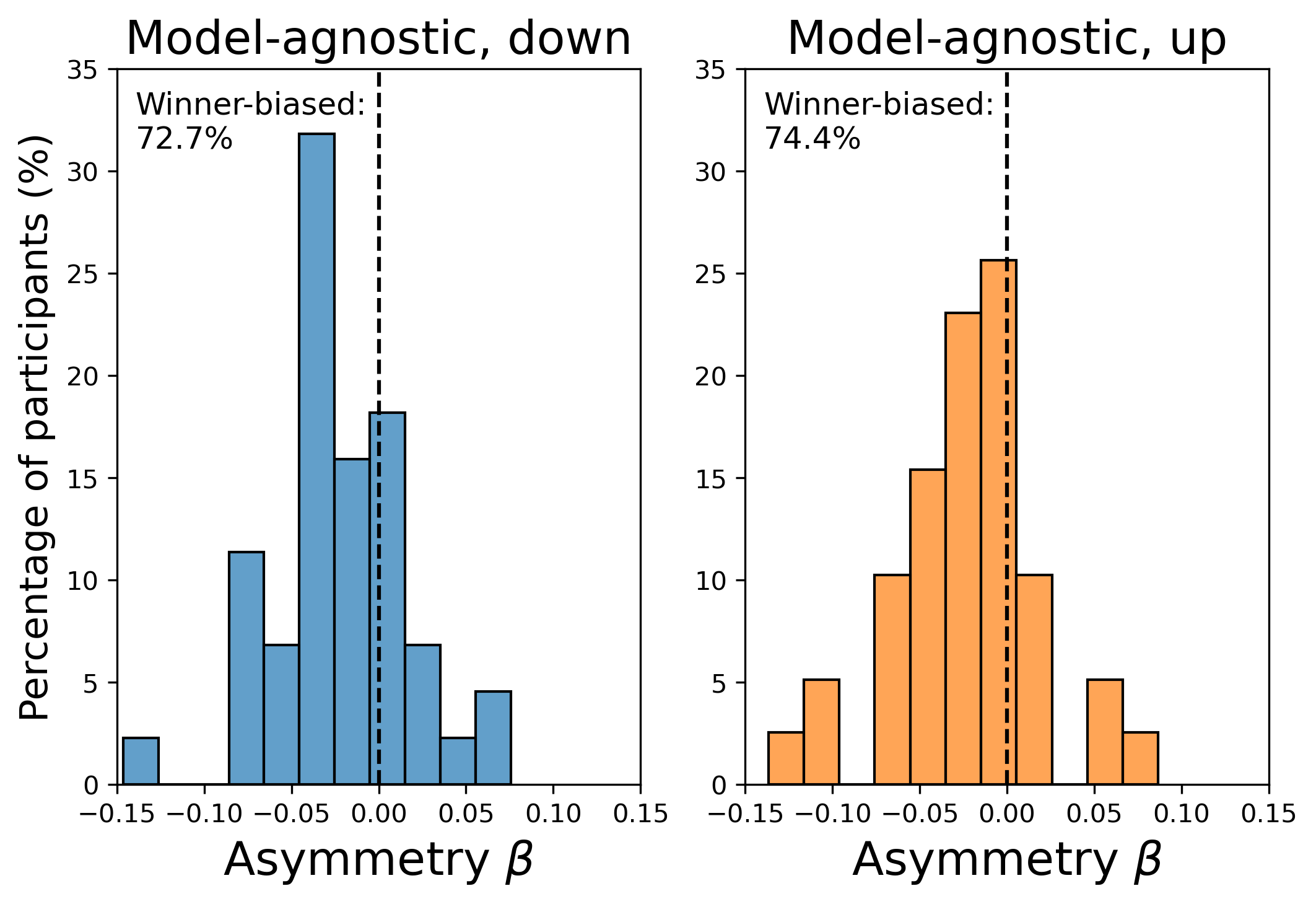

### Fig_S3.png

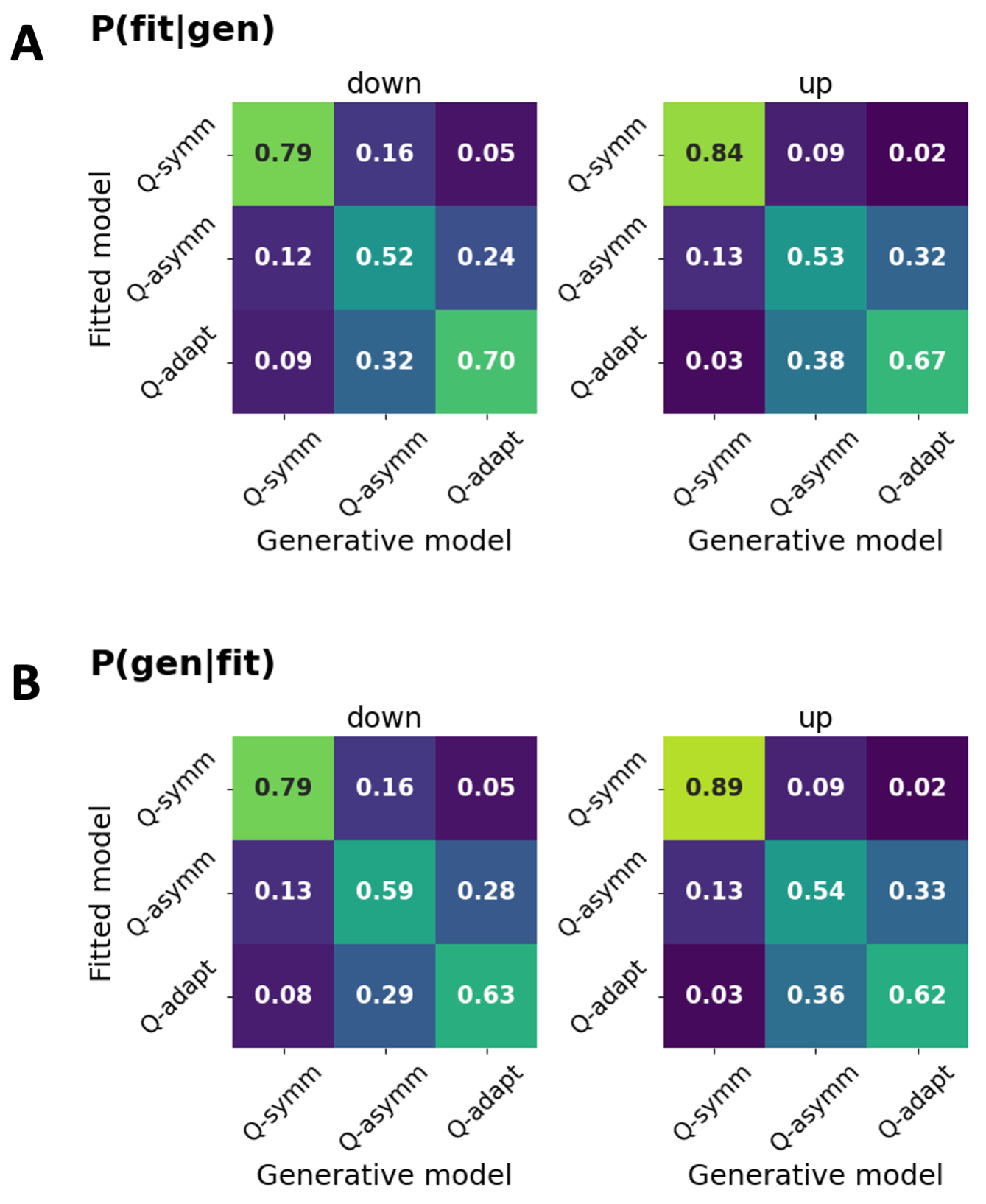

### Fig_S4.png

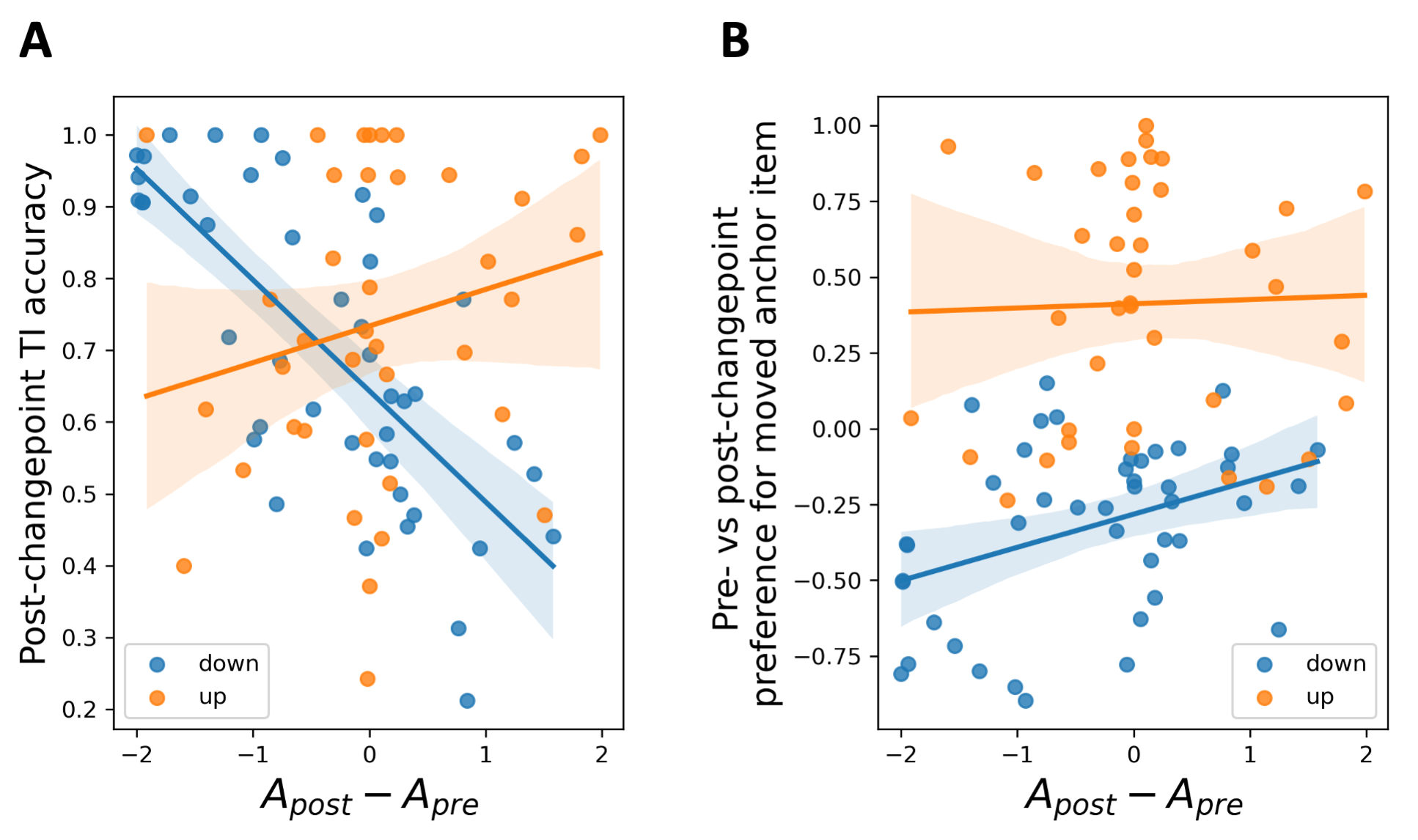

### Fig_S5.png

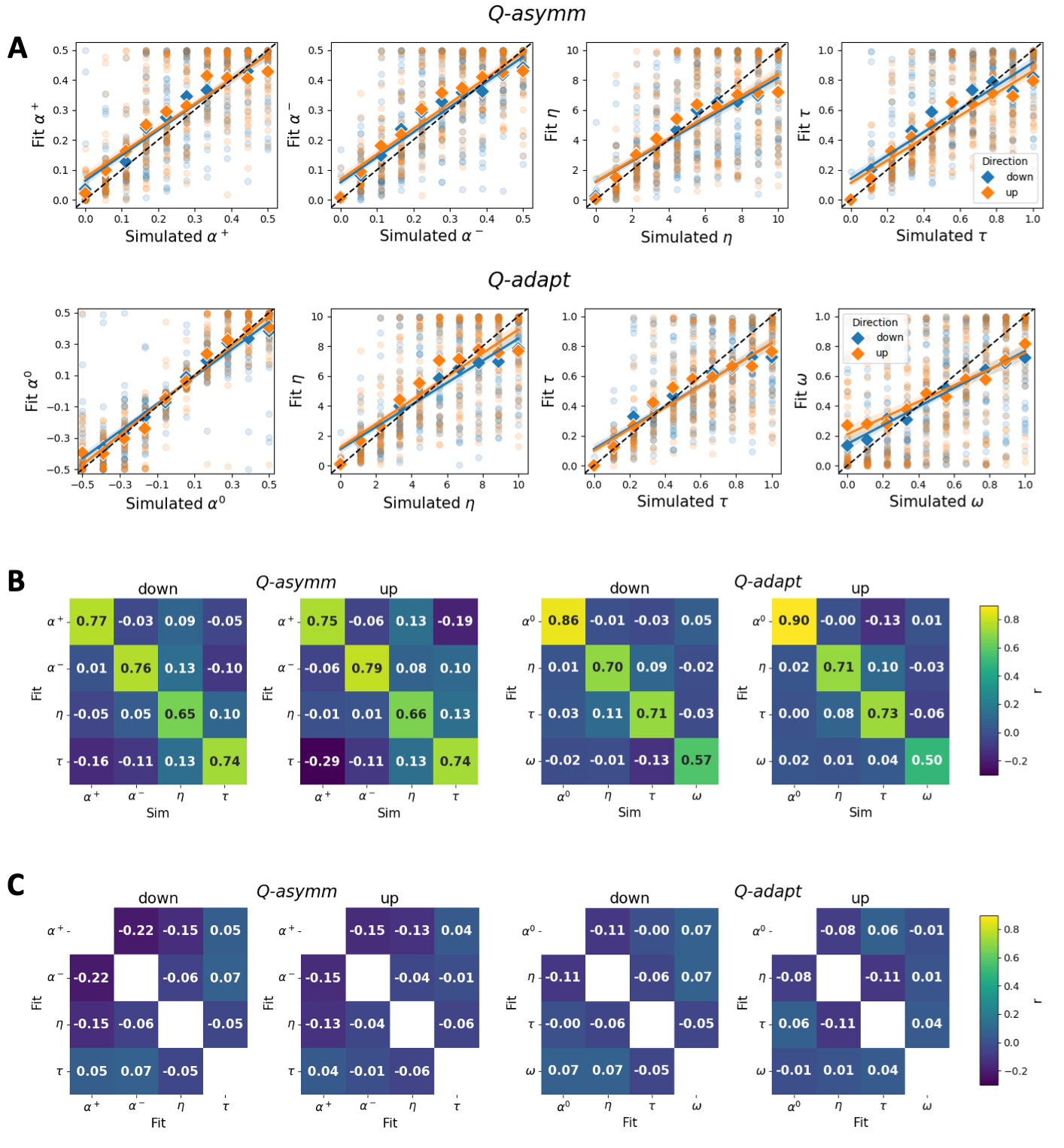
